# Theory of multiscale epithelial mechanics under stretch: from active gels to vertex models

**DOI:** 10.1101/2025.03.23.644792

**Authors:** Adam Ouzeri, Sohan Kale, Nimesh Chahare, Alejandro Torres-Sanchez, Daniel Santos-Olivan, Xavier Trepat, Marino Arroyo

**Author notes:** Sorbonne Université, CNRS, IBPS, Department of Computational, Quantitative and Synthetic Biology (CQSB), Paris, France. Department of Mechanical Engineering, Virginia Tech, Blacksburg VA, USA.

## Abstract

Epithelial monolayers perform a variety of mechanical functions, which include maintaining a cohesive barrier or developing 3D shapes, while undergoing stretches over a wide range of magnitudes and loading rates. To perform these functions, they rely on a hierarchical organization, which spans molecules, cytoskeletal networks, adhesion complexes and junctional networks up to the tissue scale. While the molecular understanding and ability to manipulate cytoskeletal components within cells is rapidly increasing, how these components integrate to control tissue mechanics is far less understood, partly due to the disconnect between theoretical models of sub-cellular dynamics and those at a tissue scale. To fill this gap, here we propose a formalism bridging active-gel models of the actomyosin cortex and 3D vertex-like models at a tissue scale. We show that this unified framework recapitulates a number of seemingly disconnected epithelial time-dependent phenomenologies, including stress relaxation following stretch/unstretch maneuvers, active flattening after buckling, or nonreciprocal and non-affine pulsatile contractions. We further analyze tissue dynamics probed by a novel experimental setup operating in a pressure-controlled ensemble. Overall, the proposed framework systematically connects sub-cellular cortical dynamics and tissue mechanics, and ties a variety of epithelial phenomenologies to a common sub-cellular origin.

## 1 Introduction

Many of the functions of epithelial monolayers are mechanical in nature. These tissues need to maintain barrier integrity during stretch, which occurs over a wide range of time-scales and magnitudes during physiology and development. For instance, alveoli in the lungs undergo stretch cycles of more than 10 % in areal strain at breathing frequency [1]. During blastocyst hatching in the early development of mammals, the trophectoderm can increase its area by several fold within a few hours largely due to cellular stretch [2]. To avoid excessive tension upon stretch, which may compromise cell-cell junctions and lead to mechanical failure, epithelial monolayers resort to sub- and supra-cellular tension buffering mechanisms at various time-scales [3, 4, 5, 6]. Tissue tension is required for tissue-scale pulsatile contractions during dorsal closure in Drosophila, when cells change their apical area by over 60% within a few minutes [7]. Besides stretching, epithelial monolayers undergo in-plane compression caused by externally applied or actively-generated lateral confinement, which may lead to out-of-plane buckling and epithelial morphogenesis [8, 9]. Tissue shaping in 3D can also be the result of apico-basal tensional asymmetries [10, 11].

Despite significant progress [12], the quantitative characterization of the mechanics of epithelial sheets *in vivo* remains very challenging, and only a few *in vitro* techniques have been developed to measure the mechanical response of cell monolayers to stretch or compression. Free-standing cell monolayers have been tested in uniaxial creep or stress relaxation by suspending them between rods [13, 14]. Tissue tension has also been measured using traction force microscopy (TFM) during stretch/unstretch maneuvers in cell monolayers adhered to deformable substrates [15]. Finally, tissue tension has been mapped using free-standing and pressurized epithelial sheets of controlled shape (epithelial domes), reaching areal strains of up to 300% [4, 16]. These tests have identified a wealth of epithelial time-dependent phenomenologies including reinforcement [15] and varying resting-length elasticity [3] under stretch, fluidization [15] and transient buckling [17] under compression, fluid or solid rheology under creep [13], tissue curling [18], and active superelasticity [4]. These approaches enable the mechanical characterization of a living tissue, while performing live imaging of structural proteins as well as biochemical perturbations, and hence they provide an experimental method to connect sub-cellular dynamics and tissue mechanics. Focusing on time-scales shorter than those required for junctional network remodeling, which depending on the jamming state of the system can range from tens of minutes to hours, these studies have shown that many of these tissue behaviors are controlled by the dynamics of the actomyosin cortex [4, 3], although at extreme deformations the intermediate filament cytoskeleton becomes mechanically active and plays an important role [4, 19].

In contrast to these experimental advancements, conventional vertex models used to theoretically study epithelial sheets build on the capillary theory of foams, and implement dynamics through either generic drag forces [20, 21, 22, 23, 24, 25] or the insertion of *ad hoc* rheological elements [26]. A theoretical and computational framework connecting epithelial time-dependent mechanics to the physics of cytoskeletal and adhesive complexes at a sub-cellular scale has been lacking [27, 28]. Such framework could help elucidate how mechanisms within cells determine the emergent and time-dependent mechanics of tissues. As an initial step, here we develop a theoretical and computational framework that upscales actomyosin cortical dynamics within cells to the epithelial scale, assuming that the junctional network does not change. To describe the out-of-equilibrium dynamics of the actomyosin cytoskeleton, we resort to active gel theories [29], modeling active force generation, remodeling of the transient network, and molecular turnover. These models have been used to understand how the actin cytoskeleton controls the mechanics of single cells [30, 31, 32, 33], and have the potential to incorporate the coupled dynamics of multiple species including signaling molecules [34] and order parameters describing the network architecture [35]. Here, we put forth a vertex model for epithelial tissues, whose time-dependent mechanics result from the net effect of an ensemble of active gel surfaces enclosing cellular volumes. In a steady state, this active-gel vertex model (agVM) recovers classical 3D vertex models [8, 22, 11]. Out of equilibrium, however, it differs from conventional dynamical vertex models in that tissue mechanics now results from cytoskeletal dynamics.

After presenting the active gel model used here and the procedure to assemble ensembles of active gel surfaces into a tissue, we show how this approach recapitulates, within a single framework, all the time-dependent phenomenologies discussed in the second paragraph, including some in which tissue behavior does not parallel single-cell behavior due to non-affinity. Furthermore, we apply the proposed computational model to study a novel experimental approach implementing a pressure-controlled mechanical ensemble representative of pressurized epithelia. Finally, we demonstrate how our multiscale modeling approach is able to predict non-reciprocal pulsatile modes similar to those observed in vivo, suggesting a built-in mechanism in epithelial tissues supporting the control of tissue tension control and locomotion.

## 2 Theoretical and computational model

### 2.1 Active gel model of the actin cortex

The actin cortex, a thin layer of contractile actomyosin gel adjacent to the plasma membrane [36, 37, 38], is composed of cross-linked actin filaments, and thus it can behave as an elastic network of semi-flexible filaments. This behavior, however, is restricted to short time-scales, between a few seconds to about a minute, because all components of the network undergo a continuous cycle of assembly and disassembly. Unbinding of a cross-linker or filament disassembly locally release network elastic energy, which is dissipated in the frictional environment. In turn, the next cross-linker binds a partially relaxed network. Thus, at longer time-scales, the cortex flows as a viscoelastic fluid [39]. Another key aspect of the cell cortex is the transduction of chemical energy into mechanical work through myosin motors or the out-of-equilibrium actin polymerization and binding of cross-linkers, effectively resulting in a contractile active tension dependent on network architecture [37].

Theoretical models of the actin cortex necessarily involve a drastic coarse-graining of the real system, in which over 100 proteins build and regulate the network [38], but still capture its fundamental physics at various scales. Below the cellular scale, microscopic models of individual filaments, cross-linkers and motors can dissect the relation between network connectivity and contractility [40, 41]. At cellular scales, active gel theories provide mechanistic understanding of cell mechanics, morphogenesis and motility using a continuum hydrodynamical description [42, 30, 31, 32, 33]. Here, we consider a relatively simple isotropic active gel model for an active viscoelastic medium undergoing turnover in the spirit of [43], but the upscaling framework presented here is transparent to the specific choice of active gel model for the actin cortex and is thus compatible with further refinements of the theory.

We summarize next the active gel model for cortical surfaces, and refer to Appendix A for a detailed derivation. We view the cortex as a quasi-2D layer defining a surface. The shape of a cortical patch, e.g. a cellular face, is given by a time-dependent surface parametrization *φ* : Γ_0_×(0, *T*) → ℝ^3^, which at each instant *t*∈ (0, *T*) maps an arbitrary planar reference configuration of the cortical surface Γ_0_ to a deformed surface Γ_*t*_ = *φ*(Γ_0_, *t*). We describe Γ_0_ and the ambient space with Cartesian coordinates, *X*^1^, *X*^2^ and *x*^1^, *x*^2^, *x*^3^. The Lagrangian velocity field is given by ***V***(***X***, *t*) = ∂_*t*_*φ*(***X***, *t*). The in-plane deformation of the cortical surface is characterized by the right Cauchy-Green deformation tensor 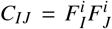, where 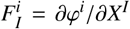 for *I* = 1, 2 and *i* = 1, 2, 3 is the deformation gradient. The Jacobian determinant of the transformation 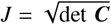, measures local area changes.

We define on Γ_*t*_ a field of areal density of cortical material *ρ*. If the actomyosin gel has constant volumetric density, then *ρ* is proportional to the cortex thickness. Likewise, we can define a field of nominal density per unit reference area in Γ_0_, denoted by *ϱ*_0_ = *J*_*ρ*_. Mass balance of the cortical material undergoing turnover is expressed by

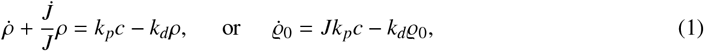

where the dot denotes material time differentiation, that is time differentiation at fixed ***X***, *k*_*p*_ and *k*_*d*_ are polymerization and depolymerization rates, and *c* is the cytosolic concentration of cortical materials ready for polymerization [4]. Assuming *c* = *c*_0_ remains constant, the steady-state cortical density is 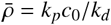.

At short times, the cortical network behaves like a nonlinearly elastic 2D material described by a hyperelastic energy density per unit mass Ψ(***C***), which if isotropic, should depend on two invariants of the tensor ***C***, e.g. *I*_1_ = trace ***C*** and *I*_3_ = det ***C*** [44, 45]. The calculation of these invariants implicitly involves the metric tensor of Γ_0_, which for our Cartesian coordinates is the identity. To represent network remodeling caused by the transient nature of cross-linkers and filaments, we introduce a material metric tensor, whose contravariant components are denoted by ***G***^***IJ***^**(*X***, *t*), and view it as a dynamical variable describing the change in “resting strain” as the system undergoes dissipative internal relaxation [46, 47, 48, 49]. We thus define the modified invariants relative to this material metric as

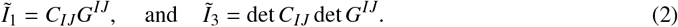

This new dynamical field allows the system to reach an elastically relaxed state irrespective of the magnitude of the deformation. Indeed, when *G*^*IJ*^ = *C*^−1,*IJ*^ we have 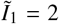 and 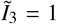, which naturally corresponds to the minimum of Ψ(***C, G***). This state can be physically interpreted as one in which the current configuration has become the reference configuration to measure elastic deformations. This approach can be interpreted as the multidimensional generalization of a 1D spring element with variable resting length [3], which here is a second-order symmetric tensor field.

The total (2D) stress tensor of the cortical layer has an elastic component, that depends on the stored energy function Ψ and that relaxes over time, and an active isotropic part, which we assume to be proportional to cortical density. The elastic part of the stress, expressed in the Lagrangian frame as the second Piola-Kirchhoff stress tensor, can be computed as ***S***^e^ = 2*ϱ*_0_∂Ψ/∂***C***. We denote the Cauchy stress tensor, the Eulerian counterpart of ***S***^e^, with the symbol *γ* and reserve *σ* for the 2D stress (surface tension) of the tissue monolayer. Following conventional relations in continuum mechanics, the elastic Cauchy stress tensor is *γ*^e^ = (1/*J*)***FS***^e^***F***^*T*^. The total Cauchy stress tensor of the surface, accounting for active tension, then takes the form *γ* = *γ*^e^ + *ξ*_*ρ*_ ***g***, where *ξ* is an activity parameter and ***g*** is the metric tensor of Γ_*t*_. We denote by 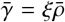 a reference active tension at the steady-state density. Force balance on the cortical surface in a low Reynold’s number regime can be expressed in terms of a tangential component

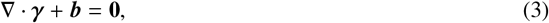

where ∇· is the surface divergence, and a normal component

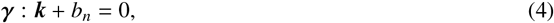

where ***k*** is the extrinsic curvature tensor of the surface. The tangential force density ***b*** and its normal counterpart *b*_*n*_ account for the tractions resulting from the interaction with the surrounding cytosolic or extracellular media.

The dissipative evolution law for the material metric is driven by the viscoelastic stress, tends to relax the stored elastic energy, and takes the form

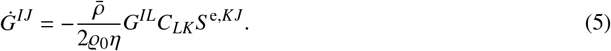

where *η* is a viscosity parameter. This equation, along with mass balance and linear momentum balance in Eqs. (1,3,4), govern the dynamics of shape and flow, given by ***V***, of density *ρ* and of material metric ***G*** of the active viscoelastic gel. The resulting model corresponds to an active viscoelastic fluid, similar to classical ones [29] but with a hyperelastic response at short time-scales and hence compatible with nonlinear models for networks of semi-flexible filaments [50]. Furthermore, having a hyperelastic limit guarantees by construction thermodynamical consistency in the sense that elastic work is zero in closed deformation paths at fixed ***G*** and *ρ*, and hence elasticity does not contribute to power input or dissipation. This is not the case in general for models having hypoelastic limits [51]. For simplicity, here we take Ψ to be a 2D-compressible neo-Hookean hyperelastic model characterized by Lamé parameters *λ* and *µ*, which we consider as integrated through the cortex thickness, and hence have units of surface tension.

Several time-scales emerge from this model, including the turnover time *τ*_to_ = 1/*k*_*d*_, the viscoelastic time *τ*_ve_ = *η*/*λ*, and the viscoactive time 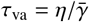. The two last time-scales are related through the dimensionless number 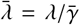, measuring the relative weight of cortical elastic modulus and active tension, which can therefore be interpreted as an elastocapillary number.

### 2.2 Building a tissue from sub-cellular active gels

We place ourselves in a regime in which tissue deformation is accommodated by changes in cellular shape rather than topological changes in the junctional network, and assume that tissue mechanics is controlled by the mechanics of cellular surfaces. The negative tension caused by adhesion molecules in cell-cell junctions contributes to the total tension of these surfaces [52]. Here, we do not explicitly model this effect, implicitly assuming that it can be subsumed in the active gel model. However, our formalism can easily accommodate more detailed models of adhesion dynamics [53, 27]. We further assume that cells keep their volume fixed, but models of cell volume regulation could be included [54, 55].

As a result of these assumptions, our model can be summarized as an ensemble of active gel surfaces enclosing 3D regions of constant volume. Computationally, we solve the fully nonlinear partial differential equations of the cell cortex of each individual cell in the tissue, Fig. 1a,b, governing the hydrodynamics and shape dynamics of the thin gel accounting for its viscoelastic rheology, active force generation and turnover dynamics. These equations are approximated by surface finite elements [56], Fig. 1c. In this approach, the discrete unknowns are the velocity and cortical density at each finite element node, and the reference metric tensor in each triangle. For computational efficiency, we assume that *ρ* and ***G*** are constant over cell faces as we do not focus on sub-cellular cortical flows driven by contractility gradients. We refer to this modeling procedure as the active gel Vertex Model (agVM) in the remainder of the paper. The agVM can be further coarse-grained to a degree in which each cell is a polyhedron whose configuration is given by the position of its vertices, Fig. 1d, which we refer to as the coarse agVM, (CagVM). At steady state our active gel model results in constant surface tensions at cell surfaces given by 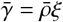, which are balanced by cellular pressures *P*_*c*_ keeping volume fixed, Fig. 1e. Thus, at steady state, the agVM is a 3D vertex model with curved surfaces [57, 58, 11, 25, 59], sometimes called bubbly vertex models, whereas the CagVM results in a conventional 3D vertex model [22].

**Figure 1:**
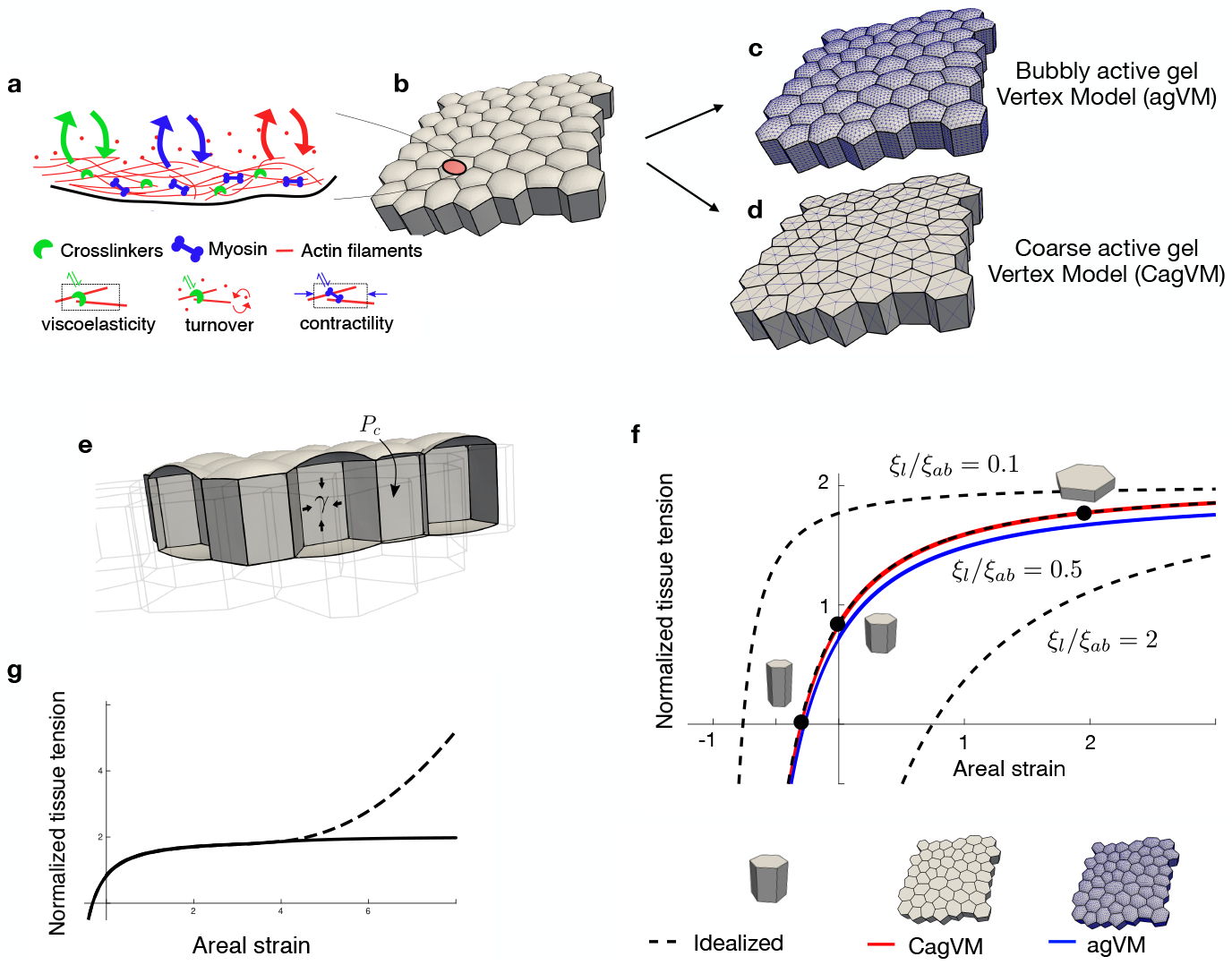
The active gel Vertex Model (agVM) and tension-strain relation at steady-state. To connect cytoskeletal dynamics within each cell and tissue mechanics (a,b), the governing equations for active gel dynamical surfaces are solved using surface finite elements in the agVM. (c) If the finite element mesh is sufficiently fine, then the result is a bubbly active gel Vertex Model, which we refer to as agVM. (d) Coarse agVM (CagVM) based on a coarse mesh whose shape depends only on the vertices of polyhedral cells. (e) At steady state, agVM and CagVM are conventional vertex models with constant surface tensions on each apical, basal and lateral face, and with cellular pressures enforcing constant cell volume constraints. (f) Normalized tissue tension, 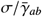 as a function of areal strain predicted by the agVM, the CagVM, and an idealized tissue of prismatic identical cells, for various ratios of lateral to apico-basal surface tensions. The reference configuration of the tissue is chosen such that cells are cuboidal with a height twice the average apico-basal edge length. (g) Illustration of the relation between tissue tension and strain when the strain-stiffening elastic potential modeling the engagement of intermediate filaments above a threshold is considered (dashed line). See Table 1 for material parameters.

**Table 1:** Model parameters for Figures 1 and 2e.

| Panel | $\bar{\lambda}$ | $\lambda/\mu$ | $\tau_{ve}/\tau_{to}$ | $\xi_l/\xi_{ab}$ | $K_{high}/\lambda$ | $K_{low}/\lambda$ | $J_{high}$ | $J_{low}$ | $V_0 k_d/(S_0 k_p)$ |
| --- | --- | --- | --- | --- | --- | --- | --- | --- | --- |
| Fig. 1f | 1 | 1 | 0.5 | 0.5 | - | - | - | - | - |
| Fig. 1g | 1 | 1 | 0.5 | 0.46 | 0.033 | 166.67 | 5 | 0.12 | - |
| Fig. 2e | 0.2, 0.4, 1 | 1 | 0.5 | 0.46 | 0.033 | 166.67 | 6 | 0.12 | 10.07 |

We allow for material parameters, and in particular cortical activity, to be different at apical, basal and lateral faces, reflecting the effect of adhesionsimplemented the power-law rheology in a uniaxially stretched idealize don surface tension [52] and possibly differences in cytoskeletal dynamics due to apico-basal polarization [18]. For this, we consider different apical, basal, and lateral activity parameters, *ξ*_*a*_, *ξ*_*b*_ and *ξ*_*l*_. We define *ξ*_*ab*_ = (*ξ*_*a*_ + *ξ*_*b*_)/2 and in most calculations, unless otherwise stated, we assume that *ξ*_*a*_ = *ξ*_*b*_ = *ξ*_*ab*_. We also define the corresponding steady-state active tensions 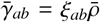 and 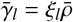.

### 2.3 Tension-strain relation of the tissue at steady-state

As a background to understand the dynamical response of cell monolayers under stretch, we first consider the steady-state mechanical response of a tissue patch by tracking the effective surface tension of the tissue *σ* as a function of equibiaxial stretch. The tissue tension *σ* collects the net effect of all cellular surfaces on the mechanics of the tissue. It is the work-conjugate to the areal strain of the tissue, defined as *ε*_*A*_ = (*A* − *A*_0_)/*A*_0_, where *A* is the actual area of the tissue and *A*_0_ is the area of the tissue in an arbitrary but fixed reference configuration. *ε*_*A*_ can be interpreted as a measure of the aspect ratio of cells relative to this reference.

In the idealized situation of a tissue with crystalline arrangement of identical prismatic cells with hexagonal apical and basal surfaces, the relation between strain and tissue tension at steady-state can be explicitly obtained as the following nonlinear relation [4]

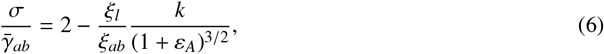

where *k* is a fixed dimensionless number, see Appendix B. Thus, even though at steady-state all elastic stresses in the actin cortex have dissipated and hence the cortex behaves like an active fluid, the assembly of tense interfaces enclosing fixed volumes behaves like a nonlinear elastic solid. The first term accounts for the contribution of apical and basal tensions to tissue tension, is independent of strain, and determines the plateau at large areal strains. The second term in this equation captures the role of lateral surfaces on tissue mechanics, and depends on strain, i.e. on cell aspect ratio. Depending on the ratio *ξ*_*l*_/*ξ*_*ab*_, the tissue tension-strain relation has different shape with different tension-free aspect ratio, Fig. 1f. In all cases, however, for sufficiently large lateral compression, i.e. for sufficiently columnar configurations, the net tissue tension becomes negative. This explicit equation for a crystalline tissue agrees very well with the response of the agVM and CagVM for a disordered tissue, from which we conclude that the details of tissue geometry are not important to predict this relation.

These results show that the maximum tissue tension at steady-state in the agVM is 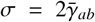 and entirely due to tension in the actomyosin cortical surfaces. However, other structures can contribute to the mechanical response of cells, including other cytoskeletal fibers, the plasma membrane under tension, or the nucleus under compression. Imaging, laser ablations, and biochemical perturbations support the mechanical role of intermediate filaments (IF) under severe stretch [13, 4, 19]. To account for this effect contributing to the re-stiffening of stretched cells, we noted the much slower turnover time of IFs as compared to actin, as well as computational work pointing at the existence of an activation strain for IF engagement [60]. Accordingly, we introduce their effect as an elastic potential activated beyond a threshold strain, Appendix C, although the strain-stiffening response of IFs is itself time-dependent [19]. The strain-stiffening regime following the tensional plateau induced by this modeling of IFs is illustrated in Fig. 1g. In the studies presented next, we introduce this effect when required as explicitly mentioned.

## 3 Results

### 3.1 Tissue dynamics under stretch

#### Equibiaxial stress relaxation

To understand how the tissue dynamically reaches the mechanical steady-state shown in Fig. 1f after a mechanical perturbation, we first turned to the experiments performed in [15] probing tissue rheology under stretch. In these experiments, Madin Darby Canine Kidney (MDCK) cells forming epithelial islands were seeded on a deformable substrate and subjected to equibiaxial stretch-unstretch maneuvers while tissue tension was quantified using monolayer stress microscopy. These experiments revealed that cell-substrate tractions were transmitted at the periphery of the cell island and that tension was nearly uniform in the central part of the tissue. Upon application of about 10% areal strain, tissue tension increased from a baseline tension to more than twice, and then partially relaxed over a few minutes to a value slightly higher than the baseline tension. Upon stretch cessation, tension decreased and then recovered the baseline over a few minutes, Fig. 2b.

**Figure 2:**
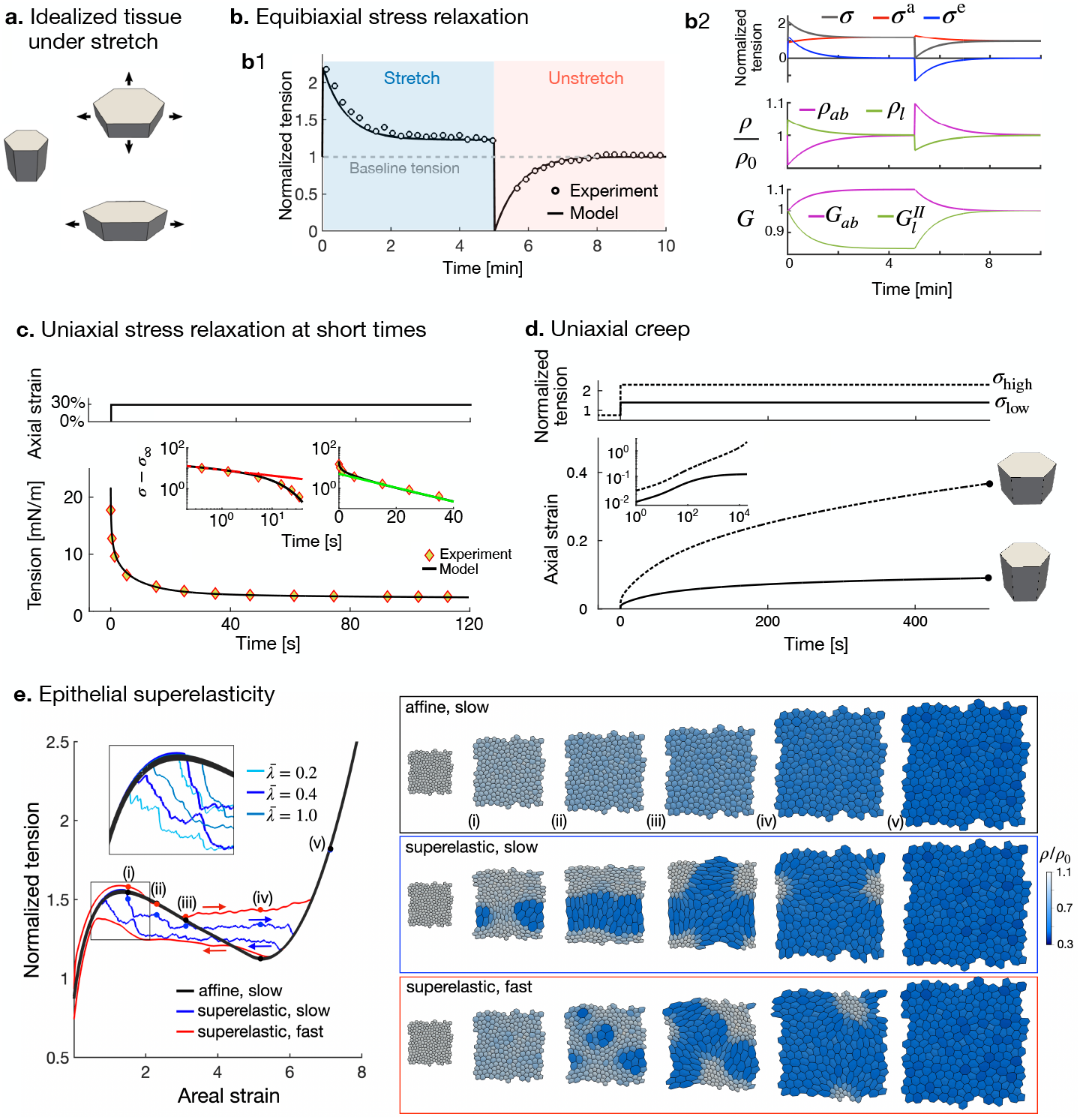
Tissue dynamics under stretch. (a) Illustration of equibiaxial and uniaxial deformations of a cell in an idealized tissue. (b1) Time evolution of tissue tension normalized by baseline tension during an equibiaxial stretch-unstretch maneuver with 10% areal strain amplitude, in quantitative agreement with the experiments in [15]. (b2) Top: Dynamics of tissue tension (total, active and elastic). Middle: dynamics of apico-basal (*ab*) and lateral (*l*) cortical densities. Bottom: dynamics of the apico-basal and lateral viscoelastic reference metric. ***G***_*ab*_ is isotropic, for ***G***_*l*_ we plot the component perpendicular to the tissue. (c) Stress relaxation following the sudden application of 30% uniaxial stretch in a model including the cortical active gel and a cytosolic power-law element along with the data points from [3]. The log-log and semi-log insets highlight the initial power-law relaxation with exponent *α* = -0.28 (dashed red line is power-law fit), followed by an exponential relaxation with timescale *τ* = 12.7 s *τ*_ve_ (dashed green line is exponential fit). (d) Creep dynamics for two magnitudes of applied tension, showing a solid-to-fluid transition when the applied tension exceeds 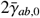. Tension is normalized by 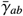. (e) Epithelial superelasticity, induced by softening due to exhaustion of cytoskeletal material and re-stiffening due to intermediate filaments, simulated with the agVM during a stretch-unstretch cycle, see also Movie 2. Fast 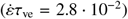 and slow 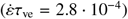 agVM simulations are compared with simulation of a tissue deforming quasistatically and constrained to remain affine. The inset in the tension-strain plot shows influence of the elastocapillary parameter 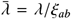 for slow agVM simulations (the reference value in the rest of the figure is 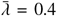). See Tables 1 and 2 for material parameters.

To test whether our model is able to recapitulate this phenomenology, we focused on the central part of the island and subjected the CagVM and agVM models to equibiaxial stretch mimicking the experiments. We further considered a simplified model considering an idealized tissue with identical hexagonal cells with planar cortical patches, Fig. 2a. This model results in 5 coupled ordinary differential-algebraic equations for the dynamics of turnover, viscoelastic relaxation and strain, see Appendix B, and therefore is extremely efficient computationally. According to this model, tissue tension can be additively decomposed into active and elastic components, *σ* = *σ*^a^ + *σ*^e^, with *σ*^e^ relaxing over time viscoelastically. For the CagVM and agVM models, tissue tension can be obtained using standard techniques of computational homogenization [61].

**Table 2:** Model parameters for Figure 2.

| Panel | $\bar{\lambda}$ | $\lambda/\mu$ | $\tau_{to}$ | $\tau_{ve}$ | $\tau_{ve}/\tau_{to}$ | $\xi_l/\xi_{ab}$ | $\alpha$ | $C_\alpha$ | $\bar{\gamma}_{ab}$ | $C_\alpha h_0/(\bar{\gamma}_{ab} \tau_{ve}^\alpha)$ |
| --- | --- | --- | --- | --- | --- | --- | --- | --- | --- | --- |
| Fig. 2b | 1 | 1 | 60 s | 45 s | 0.75 | 0.55 | - | - | - | - |
| Fig. 2c | 1 | 1 | 30 s | 10 s | 0.33 | 0.4 | 0.4 | 2534 Pa s$^\alpha$ | 0.975 mN m$^{-1}$ | 10.35 |
| Fig. 2d | 1 | 1 | - | - | 0.33 | 0.4 | 0.4 | - | - | 10.35 |

All three models provide undistinguishable results (Fig. 8a), and very accurately reproduce the experimental dynamics of tissue tension, Fig. 2b1. According to our model, the phenomenologies qualified in [15] as “reinforcement” after stretch and “fluidization” after unstretch result from cytoskeletal dynamics within cells involving changes in cortical density due to turnover and in reference metric due to viscoelastic relaxation, Fig. 2b2 and Movie 1. In our simulations, fast stretch increases/decreases the surface area of apico-basal/lateral cell surfaces, instantaneously diluting/concentrating the cortex before turnover reestablishes 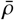 over the time-scale *τ*_to_ of about two minutes. This stretch-induced dilution/concentration of cortical material affects active tensions in cellular surfaces, resulting in a small decrease of the active component of tissue tension *σ*^a^, Fig. 2b2. The changes in shape of apical, basal and lateral surfaces following fast stretch lead to elastic cortical tensions, that result in a sudden increase of the elastic component of tissue tension *σ*^e^, which relaxes over a time-scale of *τ*_ve_ as the viscoelastic metric tensors adapt to the current strain. These dynamics are reversed upon unstretch.

To select the material parameters, we chose an initial state with cell aspect ratio comparable to the experiments (*h*_0_/*s*_0_ = 2 with *h*_0_ the height and *s*_0_ the length of an apical cell-cell contact) and selected the steady-state ratio of lateral and apico-basal surface tension 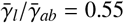 to fit the steady-state tension after stretch application. Turning to dynamics, as discussed in the previous paragraph, the active and elastic components of tissue tension have opposite trends upon stretch/unstrech. Since tissue tension increases/decreases upon stretch/unstrech, we conclude that changes in elastic tension dominate tension dynamics, and consequently “reinforcement” and “fluidization” phenomenologies of cell monolayers [15] result mostly from short-term elastic cortical tensions and their viscoelastic relaxation. We fit the viscoelastic parameters 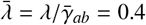 and *τ*_ve_ = *η*/*λ* = 45 s to reproduce the tension relaxation dynamics, finding a remarkable agreement with the experiments, Fig. 2b, with parameters consistent with previous estimations based on single cells, Appendix D. Although the equations controlling *σ*(*t*) are highly nonlinear, the active and the elastic components of stress after stretch and compression closely follow exponential relaxations controlled by *τ*_to_ and *τ*_ve_ respectively, Appendix B and Fig. 8b. Because viscoelastic relaxation dominates tension dynamics, tissue tension is well approximated by a single exponential relaxation typical of a standard linear solid.

#### Uniaxial stress relaxation and creep

The ability of the setup in [15] to probe the time-scales of stress dynamics below 10 or 20 s is limited by the need to refocus to acquire images of bead positions for traction force microscopy after stretch perturbation. The dynamics of stress relaxation at faster timescales can be tracked with a different setup, in which a free-standing cell monolayer is suspended between two rods enabling application/measurement of stress/strain [13]. Experiments probing the shorter time-scales have revealed a biphasic stress relaxation, with an initial ATP-independent power-law relaxation, previously identified in single cells [62], followed by an exponential relaxation [3, 63]. As shown by the previous example, our agVM describes the second of these behaviors. To examine the generality of the present modeling approach, we incorporated the power-law rheology to the tissue model. The physical origin of the required broad spectrum of relaxation mechanisms is not clear and may involve distributions of filament lengths or of cross-linker time-scales. In the absence of precise information, we introduced a spring-pot power-law element associated to bulk cytosolic/cytoskeletal shear rate, Appendix E and examined the response of the tissue under uniaxial stretch, Fig. 2c,d. This power-law element depends on two additional parameters, a power-law exponent *α* and a parameter controlling the strength of this element, *C*_*α*_.

The model predicts a biphasic power-law/exponential behavior in a stress relaxation test, Fig. 2c, which allowed us to easily fit the active gel and power-law element material coefficients to experiments reported in [3, 63]. By fitting our model to these experiments on free-standing MDCK monolayers, we found faster turnover and remodeling rates as compared to MDCK monolayers attached to a matrix, suggesting that attachment to the matrix may slow down cytoskeletal dynamics.

We then wondered if the model is able to describe the creep response as probed on free-standing monolayers under uniaxial stress. One striking experimental observation is that, while for low tensions MDCK monolayers behave like viscoelastic solids reaching a steady-state strain, for high tensions they flow like complex power-law fluids, at least within a range of deformations [13]. Our modeling framework provides a simple mechanism for this solid-fluid transition, since at steady-state it predicts a maximum tissue tension, Fig. 1f. Hence, if applied tension is above this threshold, the system cannot reach steady-state and flows, at least before intermediate filaments become loaded. To test this idea computationally, we considered step tension increases above and below this threshold, Fig. 2d, recovering the solid-like response at low tensions and the fluid-like power-law behavior when applied tension exceeds 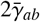 (inset).

#### Epithelial superelasticity

We attribute the agreement between the idealized tissue model, the agVM and the CagVM in the situations examined so far to the affinity of deformations, according to which individual cells experience nearly the same deformation as the overall tissue. Non-affinity is generally expected when the material is highly heterogeneous or contains unstable elements, which can result in localization of deformation. Cell monolayers under stretch have been reported to develop strong non-affinity during epithelial superelasticity [4], as a result of stretch induced cortical dilution and mechanical softening, followed by re-stiffening due to intermediate filaments. To account for the limited amount of cytoskeletal material responsible for stretch-induced softening, we fixed the total mass of actomyosin in each cell, computed as the integral of *ρ* over all cellular surfaces plus the cell volume times the cytosolic concentration *c*, Eq. (1), see Appendix B.4. To account for re-stiffening by intermediate filaments at very large stretches, we introduced an elastic potential that activates when areal strain becomes larger than a threshold, as described in Appendix C and Fig. 1g. We performed simulations with the agVM, where a periodic tissue was stretched and unstretched at slow and fast loading rates, and compared these simulations with slow simulations where the tissue was constrained to remain affine, Fig. 2e and Movie 2. The affine simulations exhibited a non-monotonic tension-strain relation reflecting the softening-stiffening behavior of individual cells. As previously shown with a much simpler model [4], the truly multicellular simulations exhibit a strong heterogeneity of cellular deformations as cells progressively transit between a low-strain and a high-strain state, resulting in a tension-strain relation for the tissue with a long plateau, reminiscent of the Maxwell construction during a phase change. Our simulations show that increased strain rate results in delayed emergence of the superelastic behavior, and a larger hysteresis upon unloading. We interpret that cortical viscosity delays the switching of cells exceeding the unstable strain.

We observed that even in the slowest simulations, the system requires a higher tension than that predicted by the Maxwell construction to develop the superelastic behavior. A close look at our simulations showed that the system transitions collectively by switching a group of neighboring cells into the high-strain phase, which then grows largely by the propagation of the boundary of the high-strain region, Fig. 2e and Movie 2. We can therefore understand the process in terms of nucleation and growth of the high-strain phase. Consistent with this view, faster loading results in multiple nucleation regions. This physical pictures suggests that there exists an effective interfacial energy between the low- and the high-strain phases. We hypothesized that this interfacial energy should be related to the cortical elasticity, which couples nonlocally the deformation of neighboring cells. Accordingly, we found that lower/higher normalized elastic modulus 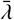 accelerates/delays the transition and results in a narrower/broader interface, Movie 2. Taken together, these results illustrate with epithelial superelasticity that non-affinity demands truly multicellular models, and suggests that the development of the superelastic transition proceeds by first nucleating domains in the high-strain phase, and by then propagating the interface, whose width and effective energy depend on 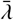.

In summary, we have shown that seemingly disparate phenomenologies in epithelial tissues, including reinforcement and fluidization under stretch/compression or superelasticity, can be explained by a theoretical model where tissue mechanics emerges from the behavior of an ensemble of active gel surfaces modeling sub-cellular cytoskeletal dynamics. Furthermore, if the tissue remains affine and isotropic and deformations nearly uniform, tissue rheology can be described with a computationally efficient model for an idealized tissue of identical prismatic cells.

### 3.2 Dynamics of epithelial domes under controlled pressure

#### The pressure-controlled mechanical ensemble with the monolayer inflator (MOLI)

Assays based on controlled in-plane stretch such as those examined in the previous section enable mechanical testing of living tissues under the canonical ensembles of imposed stress or strain. However, cell monolayers often operate in a pressure-controlled ensemble, particularly during development [64, 65]. Epithelial domes with controlled size and shape provide a natural *in vitro* system to study cell sheets under pressure [4, 16]. However, in previous implementations, pressure was self-generated by the active and polarized pumping of osmolytes. This process is transiently and stochastically disrupted by leaks, leading to uncontrolled fluctuations of pressure. To address this issue, a new microfluidic device was developed, Fig. 3a,b. This device, termed MOLI (Monolayer Inflator), was fabricated from polydimethylsiloxane (Sylgard PDMS kit, Dow Corning) as described in [66]. Briefly, the device consists of two channels, one for epithelial cell culture and the other for the application of hydrostatic pressure, separated by a porous membrane. The epithelial cell side of the porous membrane was micropatterned with fibronectin using a photopatterning technique (PRIMO, Alveole Lab), creating circular regions of 80 µm diameter with low adhesion. MDCK strain II cells expressing CIBN-GFP-CAAX were used to visualize cell membrane and tissue geometry. Upon application of hydrostatic pressure, cells on these regions detached and formed domes. The luminal pressure was controlled using a pressure channel connected to a reservoir of cell culture medium mounted on a motorized linear translation stage (Zaber), whose software enabled precise control of the fluid reservoir’s position, allowing for accurate and controlled pressure conditions throughout the experiments.

**Figure 3:**
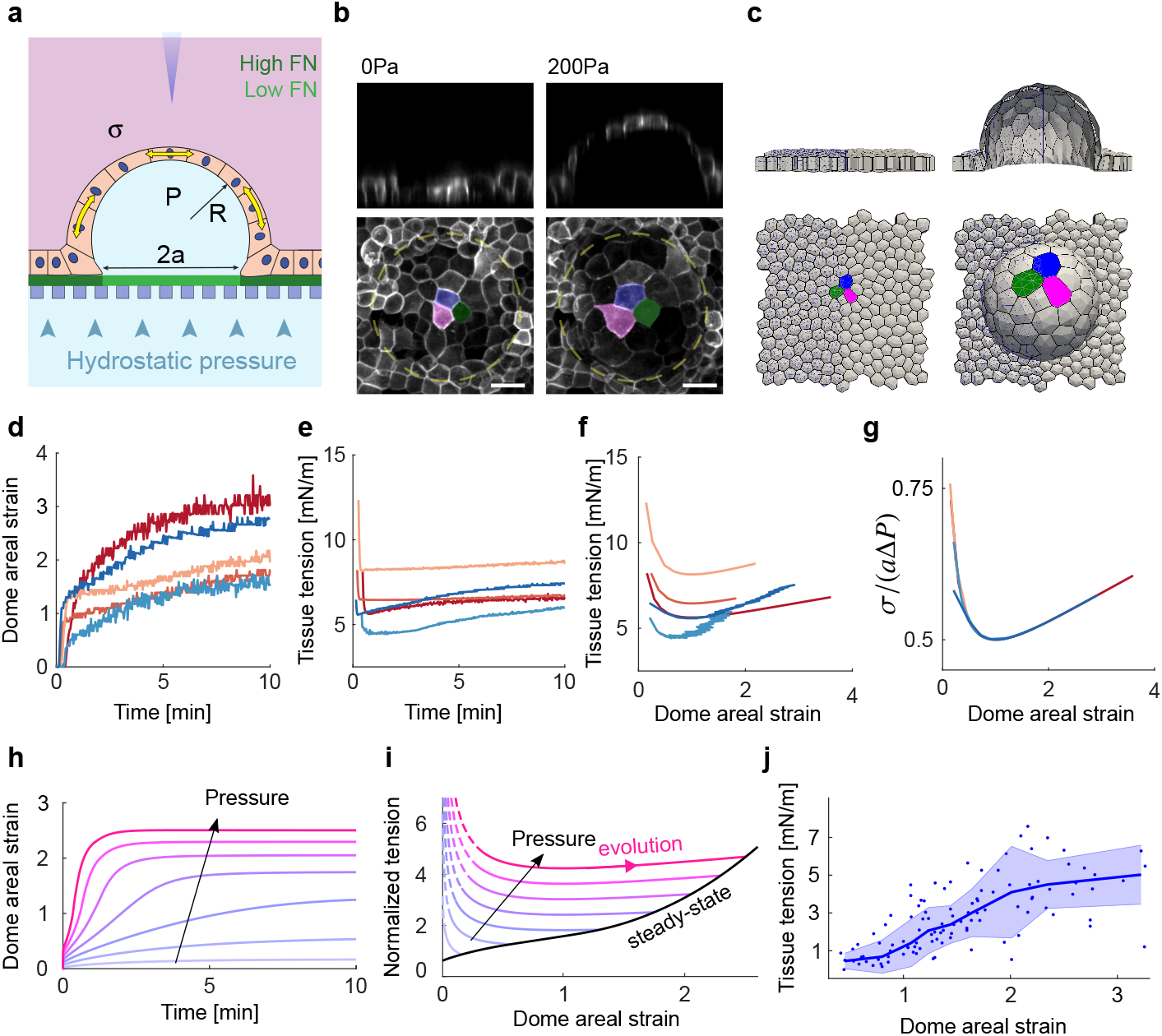
Pressure-controlled mechanical ensemble with MOLI. (a) Illustration of the experimental setup. (b) Representative confocal sections of a dome at Δ*P* = 0 and Δ*P* = 200 Pa (scalebar: 20 µm) (c) agVM simulations of the experiment. Dynamic measurements of strain (d) and tension (e) following a step pressure application of Δ*P* = 200 Pa for five representative domes with footprint radii *a* in the range 40 to 80 µm. (f) Tension-strain relation under these isobaric conditions, with data corresponding to (d,e), and (g) its normalization according to Eq. (9). (h) Strain as a function of time (normalized by *τ*_to_) for different pressure levels, applied with a fast ramp and simulated with the agVM in (c), and (i) corresponding paths in tension-strain space, converging at long times to the steady-state constitutive relation of the tissue (shown in black). (j) Steady-state constitutive relation obtained experimentally by reducing pressure stepwise and waiting in each step for 5 minutes (*n* = 12, the line and shaded area represent the median and standard deviation, with 13 points in each bin). See Table 3 for material parameters.

**Table 3:** Model parameters for Figure 3.

| Panel | $\bar{\lambda}$ | $\lambda/\mu$ | $\tau_{to}$ | $\tau_{ve}$ | $\tau_{ve}/\tau_{to}$ | $\xi_l/\xi_{ab}$ | $K_{high}/\lambda$ | $K_{low}/\lambda$ | $J_{high}$ | $J_{low}$ | $a\Delta P/\bar{\gamma}_{ab}$ |
| --- | --- | --- | --- | --- | --- | --- | --- | --- | --- | --- | --- |
| Fig. 3e,i | 1 | 1 | 38 s | 19 s | 0.5 | 0.7 | 0.333 | 166.67 | 2.3 | 0.12 | 1.211, 2.423, 3.6340,<br>4.845, 6.057, 7.268,<br>8.479 |

To understand the mechanics underlying pressure-controlled assays using MOLI, we modeled the entire dome using the agVM. To simulate the experiments, we considered an initially planar tissue, constrained the basal nodes of the model to remain in the *z* = 0 plane outside of a circular region corresponding to the region of low adhesion in the experiments, and applied a pressure difference between the basal and apical sides of the tissue, Fig. 3c. We introduced the potential restraining excessive stretch discussed earlier (Fig. 1g).

For circular footprints, the domes remained largely spherical, allowing us to simplify the analysis of tissue mechanics. Assuming a spherical cap geometry, and denoting the radius of the footprint by *a* and the height of the dome by *h*, the radius of the spherical cap can be computed as *R* = (*a*^2^ + *h*^2^)/(2*h*) and the surface area of this spherical cap as *A* = *π*(*h*^2^ + *a*^2^). Consequently, the areal strain of the dome relative to the initially planar state with area *A*_0_ = *πa*^2^ is

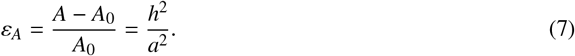

Furthermore, given the applied pressure difference Δ*P*, we can compute the tissue tension using Laplace’s law as

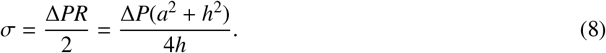

Δ*P* and *a* are experimental inputs and *h* is easily accessible at high temporal resolution by imaging single line of pixels across the midsection of the structure using line scanning mode of the Ziess LSM880 confocal microscope. Thus, these equations show that MOLI provides a method to characterize tissue rheology in a controlled pressure ensemble.

We started by applying a step pressure of Δ*P* = 200 Pa, kept it constant, and tracked strain and tension over time as shown in Fig. 3d,e. We observed that these quantities reached close to steady-states within a few minutes, consistent with the timescales of cytoskeletal dynamics and as predicted computationally by the agVM, Fig. 3h. During these experiments, we did not observe topological rearrangements of the network of cell-cell adhesions, divisions, nor extrusions. To examine the constitutive response of the tissue, we plotted tension against areal strain, Fig. 3f. Each dome exhibited a somewhat different response, but in all cases tension was very high at low strains, reached a minimum close to 100%, and then increased again at large strains. This non-monotonic response is very different to previously reported strain-tension relations for cell monolayers [13, 4, 16, 19]. To interpret it, we can combine Eqs. (7,8) to obtain

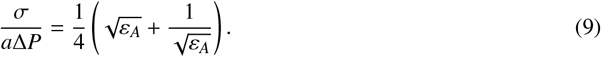

This equation shows that, given the footprint size *a* and the applied pressure Δ*P*, the dimensionless tension is proportional to 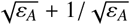 irrespective of the constitutive relation of the tissue, as this relation is a consequence of geometry and mechanical equilibrium alone. The non-monotonic nature of this isobaric tension-strain curve, with a minimum at *ε*_*A*_ = 1, reflects the fact that the radius of curvature of the spherical cap with fixed footprint is minimum when the dome is a half sphere. We represented all the isobaric tension-strain curves according to Eq. (9), finding that they collapse onto a master relation reflecting the nature of our measurement, Fig. 3g. This discussion shows that Eq. (9) alone does not provide any information about the mechanical response of the epithelial monolayer. Instead, it characterizes the locus of tension-strain states of a spherical dome with fixed footprint abiding by the physical constraint of mechanical equilibrium. It also shows the non-trivial relation between strain and stress in the pressure-controlled mechanical ensemble of MOLI. Therefore, the curves in Fig. 3d,e cannot be interpreted as creep or stress-relaxation responses, but rather as isobaric relaxation curves.

#### Quasi-static tension-strain relation

To connect our measurement with previous tension-strain data in quasi-static conditions [4, 19], we reasoned that for a given applied pressure and after a few minutes, the system should reach a steady-state characterized by a pair 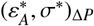 lying on the putative steady-state constitutive relation of the tissue *σ*^ss^(*ε*_*A*_). We should therefore be able to trace this constitutive relation by collecting the steady-states as Δ*P* is swept. To test this approach, we first performed simulations, where given the large strains attained experimentally we accounted for IFs as previously described, Fig. 3h,i. Experimentally, we chose to sweep Δ*P* by decreasing it, since a threshold pressure is required to initially delaminate the cell monolayer. We applied a pressure of Δ*P* = 200 Pa, waited for 5 minutes, and then decreased pressure in decrements of 20 Pa, allowing the system to relax in each step, until complete deflation. This process allowed us to collect the steady state pairs 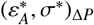, Fig. 3j, resulting in a quasi-static constitutive relation *σ*^ss^(*ε*_*A*_) that progressively increases towards a plateau of a few mN/m at high strains, in agreement with previous studies [4, 16]. In summary, these results show that MOLI can be used to map the steady-state constitutive relation of cell monolayers. They also show that this technique allows us to measure the out-of-equilibrium responses of tissues to applied luminal pressures.

#### Dynamics of epithelial domes under cyclic pressure

To examine the ability of MOLI to probe epithelial rheology out of equilibrium, we subjected domes to triangular waves of pressure with amplitude of 200 Pa and three timescales selected to be shorter, comparable and longer than the characteristic relaxation time-scale of the tissue. From confocal stacks of dome sections, we produced kymographs of dome height, Fig. 4a, which allowed us to compute the areal strain as a function of time, Fig. 4b.

**Figure 4:**
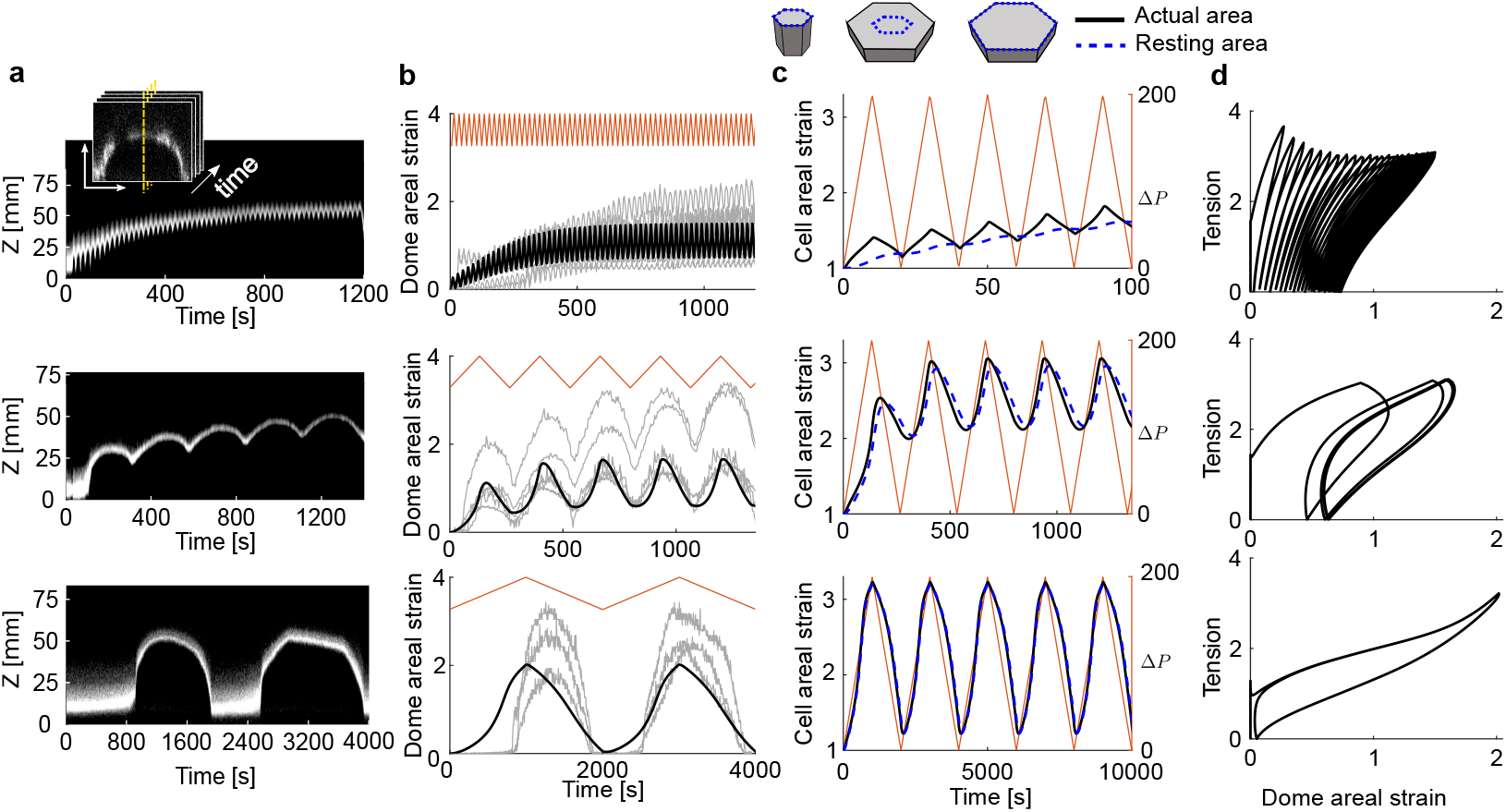
Epithelial rheology under cyclic pressure. (a) Experimental kymographs of dome height for domes subjected to cyclic pressure between 0 and 200 Pa with periods of 20 s, 266 s and 2000 s. (b) Experimentally measured areal strain as a function of time for selected domes (gray lines), and simulations with the agVM (black). The red curve indicates pressure as a function of time. (c) Dynamics of the actual area (black) and of the resting area (dashed blue) of the apical surface of a cell at the top of the dome during 5 cycles for the three different rates. The schematic at the top shows how upon sudden stretch, the resting area (computed in terms of the viscoelastic metric tensor as 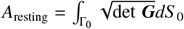) remains small and over time catches up with the actual area. (d) Dynamics of tissue tension in a dome as a function of dome areal strain according to the simulations, computed using Laplace’s law as in the experiments. See Table 4 for material parameters.

**Table 4:** Model parameters for Figure 4

| Panel | $\bar{\lambda}$ | $\lambda/\mu$ | $\tau_{to}$ | $\tau_{ve}$ | $\tau_{ve}/\tau_{to}$ | $\xi_l/\xi_{ab}$ | $K_{high}/\lambda$ | $K_{low}/\lambda$ | $J_{high}$ | $J_{low}$ | $a\Delta P/\bar{\gamma}_{ab}$ |
| --- | --- | --- | --- | --- | --- | --- | --- | --- | --- | --- | --- |
| Fig. 4b,c,d | 1 | 1 | 38 s | 19 s | 0.5 | 0.7 | 0.333 | 166.67 | 2.3 | 0.12 | 6.057 |

For the fastest cycles, the dome strain increased during pressurization and decreased during depressurization but without reaching complete strain recovery. Consequently, we observed a cumulative buildup of strain over time. After about 600 seconds (30 cycles), the maximum and minimum strains plateaued and the system seemed to reach a limit cycle. A similar response was observed for the intermediate cycles, where the system took about four cycles to reach the limit cycle. For the slowest cycles, the system fully relaxed upon depressurization to become planar, and then required a threshold pressure to detach again, presumably due to the establishment of adhesive bonds. These experiments clearly demonstrated a rate-dependent response of the domes, consistent with the viscoelastic behavior of the active gel model underlying the agVM. To further examine the ability of our model to reproduce these results, we performed simulations of cyclic pressurization, finding that with similar material parameters as in previous examples, the agVM could quantitatively reproduce the most salient features of the experiments, Fig. 4b.

To physically interpret these results, we represented over a few cycles the actual area of the apical surface of a cell at the top of the dome, along with the resting area of that surface, computed in terms of the viscoelastic metric tensor as 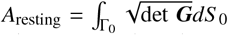 where Γ_0_ is the reference apical surface for that cell, Fig. 4c. Under the largely equibiaxial stretch of this cell, the difference between these two areas controls the elastic part of apico-basal cortical tension, which dominates tissue tension in this stretched configuration. For the fastest cycle and during the loading phase, the simulations show that the resting area significantly lags behind the actual area, resulting in positive elastic tension that contributes to resist the applied pressure and limit deformation. During the unloading phase, even though the actual area decreases, it remains larger than the resting area for a while, resulting in positive elastic tension and further increase of the resting area, see Eq. (5), which slightly decreases at the end of the cycle. This results in a net increase of the resting area over the cycle. Consequently, in the next pressurization phase, the system can deform further. This process continues in subsequent cycles, but over time, the amount of increase and decrease of resting area in a given cycle balance, resulting in a limit cycle. The limit cycle is best appreciated in the tension-strain trajectory, Fig. 4d, clearly showing that in our pressure-controlled mechanical ensemble neither tension nor strain are controlled. For cycles with intermediate rate, the behavior is similar but the difference between actual and resting area is much smaller, leading to a faster convergence to the limit cycle. In the slowest cycling rate, the resting area nearly parallels the actual area, although the out-of-equilibrium tensions are still apparent in the tension-strain trajectory, Fig. 4d.

Taken together, our agVM simulations provide a clear mechanical interpretation of experiments on epithelial domes in a controlled pressure ensemble implemented with MOLI. They also show that the tissue rheology measured under cycling luminal pressurization can be explained by the viscoelasticity of the actomyosin cytoskeleton.

### 3.3 Loss of tissue planarity

To further examine how epithelial monolayers dynamically evolve towards steady-states upon mechanical perturbations, we focused on tissue dynamics under compression. As shown in Fig. 1f, the steady-state tissue tension becomes negative for sufficiently large compression. Hence, being a slender thin film, a suspended monolayer buckles under compression as reported in [17]. This reference further reports monolayers experiencing transient buckling, followed by a recovery of planarity within a few minutes. To test whether this behavior can be explained by the agVM, we considered an initially stretched strip of tissue clamped at its ends and rapidly brought the ends closer together by various magnitudes, Fig. 5a.

**Figure 5:**
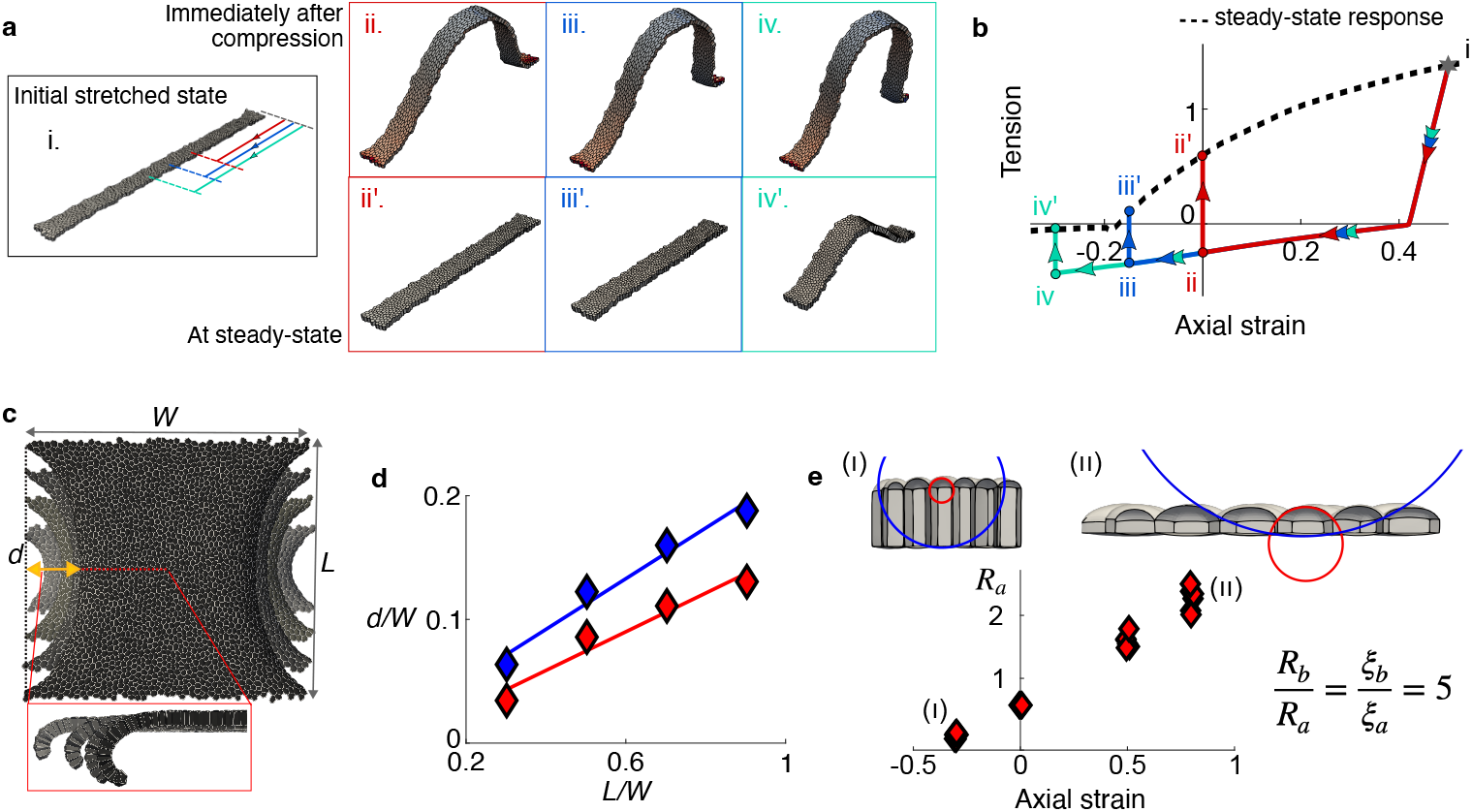
Loss of tissue planarity. (a) Rectangular tissue strip with a 50% uniaxial prestretch from a reference state under tension rapidly compressed to 0% (red), -15% (blue), and -30% (cyan) uniaxial strain, Movie 3. The snapshots show the tissue shape immediately after compression (*t* ≈ 0) and after relaxation (*t* = 10 *τ*_to_). (b) Corresponding trajectories in tension-strain space, where tension is normalized by 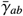. The quasistatic response of the agVM is shown with dashed black line, showing that the tissue strip is able to resist a small compressive tension at steady-state following buckling. (c) Edge curling in a tissue patch with clamped ends in the presence of apicobasal asymmetry of active tension (*ξ*_*b*_*/ξ*_*a*_ = 2) for four different tissue lengths *L*. (d) Curl deflection as a function of tissue length for *ξ*_*b*_*/ξ*_*a*_ = 1.5 (red) and *ξ*_*b*_*/ξ*_*a*_ = 2 (blue). (e) Effect of apicobasal asymmetry (here *ξ*_*b*_*/ξ*_*a*_ = 5) on the curvature of apical and basal faces. The apical radius and basal radii of curvature increases with applied strain, but according to Laplace’s law, their ratio remains constant following the relation *R*_*b*_*/R*_*a*_ = *ξ*_*b*_*/ξ*_*a*_. See Table 5 for material parameters.

**Table 5:** Model parameters for Figure 5

| Panel | $\bar{\lambda}$ | $\lambda/\mu$ | $\tau_{ve}/\tau_{to}$ | $\xi_a$ | $\xi_b$ | $\xi_l/\xi_{ab}$ | $K_{high}/\lambda$ | $K_{low}/\lambda$ | $J_{high}$ | $J_{low}$ |
| --- | --- | --- | --- | --- | --- | --- | --- | --- | --- | --- |
| Fig. 5a,b | 1 | 1 | 0.25 | - | - | 0.5 | - | - | - | - |
| Fig. 5c,d | 1 | 1 | 0.5 | 0.8, 0.6667 | 1.2, 1.3333 | 0.5 | 0.033 | 166.67 | 6 | 0.12 |
| Fig. 5e | 1 | 1 | 0.5 | 0.333 | 1.667 | 0.4 | - | - | - | - |

Based on the steady-state relation between tissue tension and uniaxial strain predicted by the idealized tissue model, we chose two compression magnitudes that should result in a positive tension at long times and a larger compression that should lead to steady-state compressive tension. In all three cases, the tissue develops negative tension as a result of the short-term elasticity of the cortex, and buckles immediately after compression, Fig. 5b and Movie 3. Over time, however, as viscoelastic relaxation and turnover take place over a time-scale of a few minutes, the two first monolayers recover planarity and positive tissue tension whereas the third monolayer remains buckled, albeit with smaller out-of-plane amplitude. Thus, in agreement with experiments [17], our model predicts that monolayers may exhibit either transient or permanent buckling depending on the degree of compression.

Besides buckling, suspended monolayers have been used to understand how an apico-basal asymmetry of active tension results in loss of tissue planarity in the form of tissue curling at the edge of free-standing monolayers of MDCK cells [18]. To interrogate curling with our model, we considered tensed tissue strips clamped at opposite ends with larger basal than apical surface tension. This introduces an active spontaneous curvature in the tissue, which readily leads to curling at the free edges of the tissue strips, with a curl morphology very similar to that in experiments, Fig. 5c. We tested the observation that the curling deflection increases linearly with tissue length [18]. We measured deflection in tissues of different lengths but equal clamped width for two different apico-basal tension ratios and recovered a linear relationship between deflection and tissue length, with the slope coefficient increasing with tensional asymmetry Fig. 5d.

Since the agVM resolves not only tissue shape but also cellular shape, we noted that in our asymmetric monolayers the apical surfaces of individual cells were much more curved than basal surfaces, Fig. 5e, in agreement with experimental observations in [18]. This fact can be easily understood from Laplace’s law, since cellular pressure is uniform in our model within each cell and hence the radius of curvature of apical and basal surfaces exhibits the same ratio as the corresponding surface tensions irrespective of the applied stretch. Pressure being stretch-dependent, our model predicts an increase in radius of curvature of apical and basal surfaces as the tissue is stretched, Fig. 5e. Thus, by connecting sub-cellular and monolayer scales, our model captures the consequences in cell and tissue shape of apico-basal polarization of active tension, enabling multiscale mechanical inference in cell monolayers [67].

### 3.4 Tissue pulsatility

Transient tissue stretch can also be the result of internal dynamics of cell monolayers in the form of tissue pulsations and traveling waves, which enable tissue morphogenesis [7, 68, 69] and multicellular migration [70, 71], and are attributed to a combination of mechano-chemical interactions in the cytoskeleton and biochemical signaling [72, 73, 43, 74, 75, 76]. 2D vertex models have been used to study such pulsations introducing phenomenological time-delays [77, 78]. Interestingly, a minimal model of the actin cytoskeleton including contractility, elasticity and turnover is sufficient to develop spontaneous oscillations reminiscent of tissue pulsations [79]. Since these features are present in the agVM through the active gel model, along with an accurate geometrical representation of the tissue, we examined whether pulsations spontaneously develop in our model and the nature of such pulsations.

The tension-strain response given by Eq. (6), Fig. 1f, collects steady-states of a dynamical system, which need not necessarily be stable across parameter space. To analyze the stability of these fixed points, we performed linear stability analysis for an idealized tissue with identical prismatic cells subjected to constant tension, Appendix B.2. For small tensions, we found that the eigenvalues of the system Jacobian matrix are real and negative, and hence the system is stable. At intermediate tensions, however, the system exhibits pairs of complex conjugate eigenvalues, whose real part becomes positive above a tension threshold, indicative of linearly unstable behavior. For even larger tensions, eigenvalues become real and the system is unstable. This picture is indicative of a Hopf bifurcation, with four regions depending on the stability of the steady states: (1) stable nodes characterized by exponential convergence from nearby states at low tension, (2) stable and (3) unstable spirals at intermediate tensions, and (4) unstable nodes with exponential divergence at high tension, Fig. 6a.

**Figure 6:**
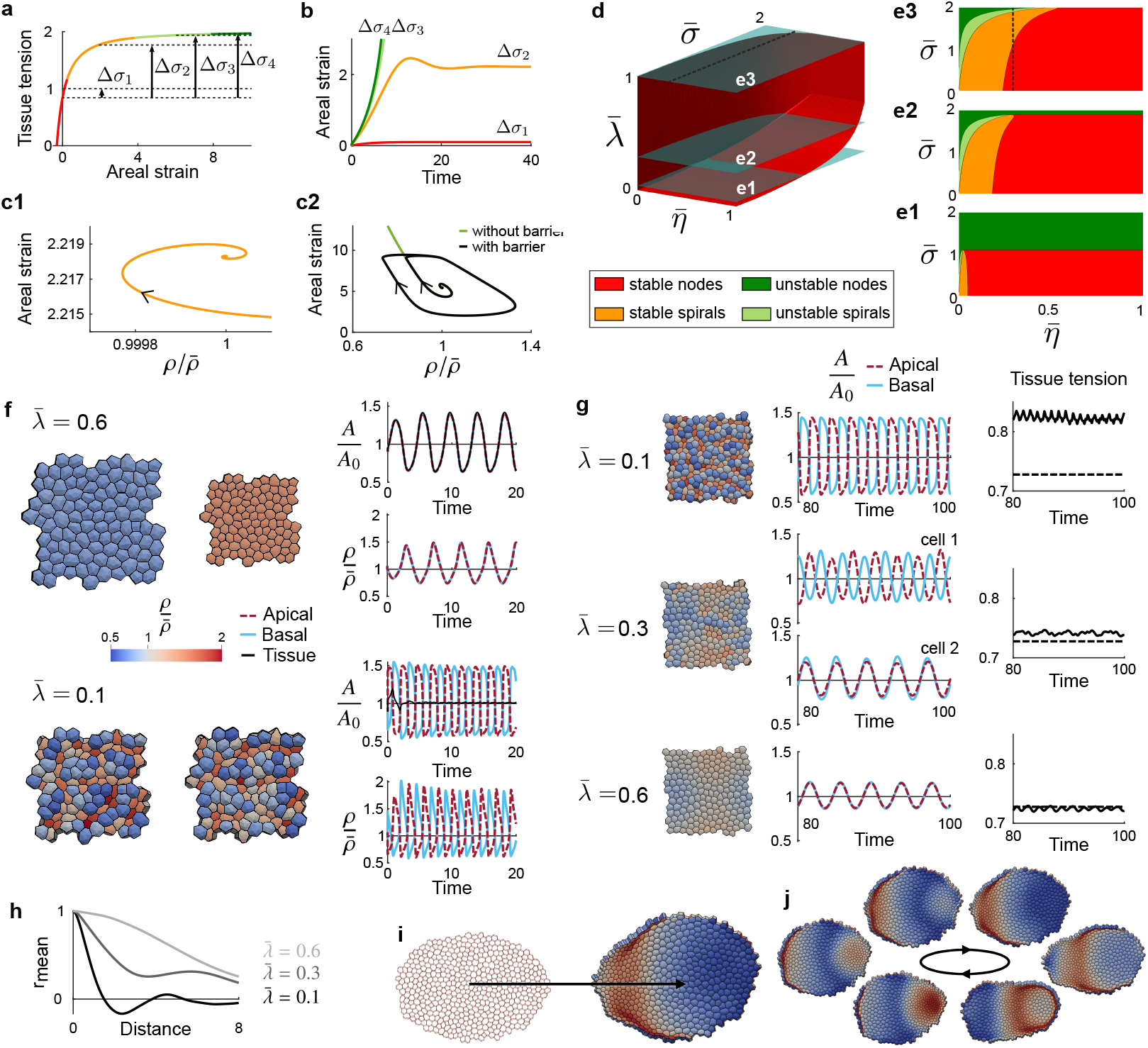
Sustained pulsating patterns modulated by the physical properties of the actomyosin cortex. (a) Steady-states in Fig. 1f color-coded depending on their dynamical stability. (b) Strain trajectories following tension increases of different magnitudes as predicted by the idealized tissue model. (c) Orbits in the density-areal strain plane after a small tension perturbation around the steady-state on either side of the stable-unstable boundary in (a). In (c2), we show the orbit of a system with a barrier limiting extreme cell deformations, leading to a stable limit cycle and sustained oscillations. (d) Stability diagram as a function of dimensionless tension, cortical elasticity and viscosity, showing the surface separating stable and unstable regions. (e) Cross-sections of the stability diagram in (d) along planes given by 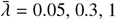. The dashed line in (d) and (e2) corresponds to (a). (f) Pulsating modes in free-standing multicellular systems at constant tension described by the agVM for two values of 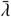. Representative out-of-phase states, and time evolution of the normalized areas (tissue, apical and basal), and of apical and basal densities. See also Movie 4. (g) Pulsating modes for a periodic tissue with constant area. Representative states, time evolution of apical and basal areas, and of average tissue tension (dashed lineas are the tension of a non-pulsating steady-state). See also Movie 5. (h) Spatial correlation averaged over time of apical density as a function of distance normalized by edge length *s*_0_, for the three values of 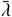 shown in (g). In all panels, time is normalized by *τ*_to_ and tension by 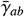. (i) Self-organized locomotion of a tissue interacting with a substrate through adhesion and friction. The arrow indicates the net displacement after 40 cycles (150 turnover times) of non-reciprocal collective pulsation. (j) Illustrative states along the non-reciprocal collective pulsation. See also Movie 6. See Table 6 for material parameters.

**Table 6:** Model parameters for Figure 6. In the last column, we report the viscous drag or friction of the monolayer with the substrate, *ζ*, in dimesionless terms as the ratio between the tissue size *L* and the hydrodynamic length 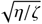.

| Panel | $\bar{\lambda}$ | $\lambda/\mu$ | $\bar{\lambda}$ | $\bar{\eta}$ | $\xi_l/\xi_{ab}$ | $K_{\text{high}}/\lambda$ | $K_{\text{low}}/\lambda$ | $J_{\text{high}}$ | $J_{\text{low}}$ | $L\sqrt{\frac{\zeta}{\eta}}$ |
| --- | --- | --- | --- | --- | --- | --- | --- | --- | --- | --- |
| Fig. 6a,b,c | 1 | 1 | 1 | 0.3 | 0.5 | - | - | - | - | - |
| Fig. 6c | 1 | 1 | 1 | 0.3 | 0.5 | 0.1667 | 0.1667 | 10 | 0.1 | - |
| Fig. 6f,g,h | 0.6, 0.3, 0.1 | 1 | 0.6, 0.3, 0.1 | 0.02 | 0.5 | 3.333 | 3.333 | 1.5 | 0.8 | - |
| Fig. 6i,j | 0.6 | 1 | 0.6 | 0.01 | 0.5 | 3.333 | 3.333 | 1.5 | 0.8 | 4.80 |

To examine the nonlinear response of the system, we integrated in time the dynamical equations of the idealized model starting from a stable node and increasing suddenly tension to values in each of these four regions. We first considered a system without the strain-stiffening mechanical barrier modeling the mechanical effect of IFs. Consistent with the linear stability analysis, the system reached stable steady states in regions (1) and (2), and diverged in regions (3) and (4), Fig. 6b. To better appreciate the dynamics of oscillatory states, we represented the orbits in the strain-apical cortical density plane, Fig. 6c, showing that the transition between regions (3) and (4) corresponds to a subcritical Hopf bifurcation diverging to infinity. By introducing the strain-stiffening behavior modeling IFs, however, we found that in region (3) the system reached limit cycles of sustained oscillations (supercritical Hopf bifurcation), Fig. 6c2.

Pulsations emerging from our model can be understood by a positive feedback between contraction-induced densification of the cortex and enhanced contractility since *γ* = *ρξ*, slowed down by viscoelasticity and the mechanical barrier, and eventually reversed by turnover, which dilutes the cortex and activates a complementary feedback loop involving cortical dilution during extension. Viscoelastic relaxation and turnover are thus built-in lag mechanisms of the actin cortex taming self-reinforcing active contraction or expansion.

In addition to tissue tension, the dynamical stability of the system should also depend on the viscoelastic parameters of the cortex. We thus systematically mapped the four regions of stability in a three-dimensional parameter space given by the key dimensionless parameters 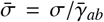 (normalized tissue tension), 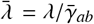 (normalized elastic modulus or elastocapillary number), and 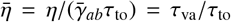 (normalized remodeling viscosity), Fig. 6d,e. We note that the steady state of the system is independent of 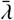 and 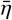. This map shows that unstable regions are associated with high tension 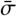 and low viscosity 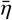, in agreement with experimental observations [69], as well as low normalized elastic modulus 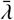. Thus, pulsatile dynamics should appear when active tension is high relative to both cortex elasticity and viscous stress buildup in the timescale of turnover.

We then examined how these pulsations of a homogeneous idealized system manifest in a truly multicellular tissue. To study pulsatile behavior, we chose initially active gel parameters leading to a non-pulsatile steady-state, and then reduced cortical elasticity to access different dynamical regimes. We first focused on a fixed tension ensemble. In agreement with the stability map above and for material parameters in the unstable spiral region, a bi-periodic agVM subjected to constant tissue tension develops a breathing mode where all cells pulsate synchronously leading to large changes of tissue area, Fig. 6f(top) and Movie 4. However, as cortical elasticity is reduced, the breathing mode becomes unstable and the system self-organizes into non-affine dynamical states not captured by the idealized theory. These states are characterized by out-of-phase pulsations of neighboring cells and of apical and basal surfaces within cells. Compensation of different cell areas results in negligible macroscopic strain changes, Fig. 6f(bottom) and Movie 4. Hence, this pulsation mode does not perform macroscopic power.

Since epithelial monolayers are generally laterally confined, we next considered a fixed strain ensemble with bi-periodic boundary conditions. For low cortical elasticity, the system develops a similar dynamical mode as in the constant tension ensemble, with out-of-phase cell-to-cell and apico-basal pulsations, Fig. 6g(top) and Movie 5. Interestingly, it results in a significant increase in average tissue tension relative to the steady-state tension in the absence of pulsations. This suggests a mechanical function for these pulsations, reminiscent of those in the amnioserosa during Drosophila Dorsal Closure [69]. At high cortical elasticity, the breathing mode of the fixed-tension ensemble is incompatible with the fixed-strain ensemble and, instead, the system develops coherent and propagating contraction waves involving tens of cells and in-phase apico-basal oscillations, Fig. 6g(bottom) and Movie 5. This dynamical mode results in negligible perturbations of the steady-state tension. For intermediate 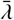, the system develops a dynamical mode combining anti-phase cell-to-cell and apico-basal pulsations with larger-scale propagating waves. These three regimes are clearly characterized by the spatial correlation of apical density capturing the lengthscale and the cell-to-cell correlation or anti-correlation of the different pulsation modes, Fig. 6h. Additional simulations with the more constrained CagVM exhibited the same phenomenology but with delayed thresholds for pulsatility, attributed to the lower spatial resolution. Thus, our results show that cortical active gel parameters control not only the threshold but also the mode of pulsation, and in particular, that cortical elasticity increases mechanical coupling between cells leading to collective pulsation modes. This role of cortical elasticity in regulating non-affinity resonates with our previous results regarding epithelial superelasticity, Fig. 2e.

A close look at the collective pulsations for large values of 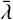, Fig. 6g(bottom) and Movie 5, shows that they are non-reciprocal, and hence may provide a self-organized mode of locomotion. To further investigate this, we considered an elliptical tissue interacting with a substrate. We assumed that the basal plane remains attached to the plane of the substrate and modeled adhesion with an effective negative free-energy proportional to basal area. Furthermore, we considered viscous drag of the basal plane with the substrate. Adhesion tends to expand the tissue, increasing 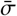 and triggering the self-organized pulsation, which are collective by our choice of 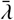. The system initially developed breathing pulsations, but after about 40 turnover times, these transitioned spontaneously to a polarized non-reciprocal limit cycle, which lead to directed locomotion of the pulsating tissue, Fig. 6i,j and Movie 6. These results are remarkably similar to the contractions during locomotion of *T. adhaerens* [70], albeit significantly slower, which we attribute to the absence of an explicit extension-induced-contraction in our model [80].

In summary, we have shown that self-organized tissue pulsations can result from the intrinsic dynamics of the actomyosin cortex modeled as an active viscoelastic gel undergoing turnover. These pulsations are controlled by three dimensionless parameters, and can be characterized by out-of-phase cell-to-cell pulsations, which enhance tissue tension, or by collective and non-reciprocal traveling contractions, which enable locomotion.

## 4 Summary and outlook

We have examined theoretically and computationally a very simple premise, that the time-dependent mechanics of a cell monolayer devoid of topological transitions [6, 4, 3, 17, 18, 7] depend on the cytoskeletal dynamics of a collection of curved cellular cortices. For this, we have developed an out-of-equilibrium framework bridging active gel models of the actomyosin cortex with vertex-like models at a tissue scale in a geometrically nonlinear setting. We have shown that a wide spectrum of epithelial phenomenologies are recapitulated by this model, including stress relaxation, creep and superelasticity under stretch, transient buckling under compression, curling of tense tissues due to apico-basal tensional asymmetry, or self-organized pulsations with different dynamical modes and different functionality. While *ad hoc* models for each of these processes have been proposed, the single model presented here describes all of them with high fidelity regarding cellular and tissue shape, further providing information about the cortical state (density and reference elastic metric). We have shown that the required spatial refinement of the active gel vertex model (agVM) depends on the nature of cellular deformations, and specifically the affinity of the tissue. In fact, our results delineate the range of applicability of continuum theories for epithelial shells, including those homogenizing vertex models [81, 82, 83, 84, 85, 86, 28, 87], which implicitly assume affinity of cellular deformations, and therefore cannot be directly applied to tissues undergoing superelastic transformations (Fig. 2e) or cell-to-cell pulsations (Fig. 6f,g).

We have assumed a very simple topology for the junctional network, but our model can be directly extended to account for scutoids [88] or other cellular arrangements. We have further assumed that the junctional network does not change and have considered a relatively simple active gel model accounting for turnover, viscoelastic relaxation of the network and active contractility. However, the framework can be adapted to tissue descriptions naturally accounting for cell rearrangements [89] and to richer multi-species models accounting for regulatory networks, and hence can provide a general background to integrate sub-cellular dynamics and emergent mechanical responses of cellular tissues. Besides the actin cytoskeleton, the framework can also accommodate other mechanical modules in epithelial tissues such as adhesion complexes [53] or volume regulation [54, 55]. As a proof of concept of this ability to combine various structural ingredients, here we have coupled the active gel description of cortical patches with a cytosolic power-law element and with an unspecific mechanical barrier limiting large strains. As opposed to minimalistic phenomenological models, our work suggests that this bottom-up modeling route has the potential to provide mechanistic understanding of active nonlinear responses of tissues and integrate high-resolution observations including cell and tissue shape, tissue mechanics, and molecular localization and dynamics.

## Acknowledgements

This work was supported by the European Research Council (CoG-681434 to MA, Adv-883739 to XT), the Spanish Ministry of Science, Innovation and Universities and the Spanish State Research Agency MICIU/AEI/10.13039/501100011033 (PID2022-142178NB-I00 to MA and PID2021-128635NB-I00 and FEDER “ERDF-EU A way of making Europe” to XT), and the Generalitat de Catalunya (2021-SGR-01049 and ICREA Academia prize for excellence in research to MA, SGR-2017-01602 to XT). IBEC is recipient of a Severo Ochoa Award of Excellence. XT acknowledges funding from the CERCA Programme and the Human Frontiers Science Program (HFSPRGP022/2024). MA and XT acknowledge the support of La Caixa Foundation (LCF/PR/HR24/00326).

## A Active viscoelastic gel description of the actomyosin cortex

We provide here a detailed derivation of the active viscoelastic gel description of the actomyosin cortex. This derivation is based on Onsager’s variational formalism to irreversible thermodynamics [90, 56, 91], which provides a direct and systematic approach to develop complex models for dissipative and active dynamics of soft biological materials. This principle is applicable when inertial forces can be neglected and in isothermal conditions, two conditions that are often met by epithelial sheets to a good approximation.

### A.1 Onsager’s variational formalism

In Onsager’s variational modeling framework, the dynamics of the system are governed by the balance of energetic, dissipative and external forces. These forces are derived from potentials, which possibly include different physics and their interactions. The system is characterized by a set of state variables *X*(*t*) and a set of process variables *V*, which describe changes in the state of the system, either simply by ∂_*t*_*X* = *V*, or more generally through a process operator such that ∂_*t*_*X* = *P*(*X*)*V*, which often encodes mass balance principles.

Given the free-energy of the system ℱ (*X*), the dissipation potential 𝒟 (*X*; *V*) and the power supplied to the system 𝒫 (*X*; *V*), we can form the Rayleighian functional, which establishes the competition between free-energy relaxation, dissipation and power input, as

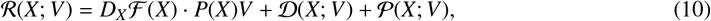

where, by the chain rule, the first term is the rate of change of free energy 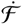. Then, the dynamics of the system are given by the instantaneous minimization of ℛ (*X*; *V*) with respect to *V*. Suppose the system is subjected to a constraint on the process variables given by ℂ (*X*)*V* = 0. Then, we can form the Lagrangian as ℒ (*X*; *V*, Λ) = ℛ (*X*; *V*) + Λ · ℂ (*X*)*V*, where Λ are the Lagrangian multipliers, and the constrained dynamics follow from

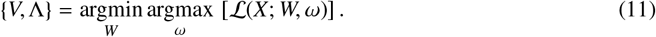

### A.2 Kinematics

We model a cortical patch as a two-dimensional surface embedded in ℝ^3^. The deformed shape of a cortical patch at time *t*, denoted by Γ_*t*_, is defined through the parametrization *φ* : Γ_0_ × (0(, *T*) → ℝ^3^, where Γ_0_ is an arbitrary planar reference configuration of the cortical patch. We denote by ***X*** = (*X*^1^, *X*^2^) points on Γ_0_ ⊂ ℝ^2^ and by ***x*** = (*x*^1^, *x*^2^, *x*^3^) points on Γ_*t*_ ⊂ ℝ^3^ described by Cartesian coordinates. As the cortical patch deforms, Γ_*t*_ evolves in time and ***x*** = *φ*(***X***, *t*) is a family of deformed configurations depending on time. In what follows, we drop the explicit dependence on time of the current surface, which we simply denote by Γ.

A parametrization *φ* induces natural basis vectors of the tangent plane of Γ given by ***g***_*I*_ = ∂*φ*/∂*X*^*I*^, where *I* ∈ {1, 2}. The inner product of these vectors defines the covariant components of the metric tensor ***g*** on Γ, *g*_*IJ*_ = ***g***_*I*_ · ***g***_*J*_, expressing the Euclidean metric of the ambient space in the surface coordinates. The contravariant components of the metric tensor *g*^*IJ*^ are defined by the relations 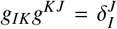. Thus, for the metric tensor, the matrix of covariant components is the inverse of the matrix of contravariant components. Likewise, we can define the metric tensor ***G***_0_ of the reference configuration Γ_0_, which is just the identity matrix when {*X*^1^, *X*^2^} are Cartesian coordinates. The natural basis vectors on Γ_0_ are denoted as ***G***_*I*_. We can define dual basis vectors ***g***^*J*^ and ***G***^*J*^ from the relations 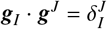 and 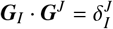. The spatial velocity is given as [44]

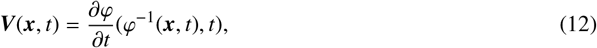

where *φ*^−1^(***x***, *t*) is the inverse of the mapping *φ*(***X***, *t*) evaluated at time *t*. The spatial velocity can be decomposed into a tangential component ***v***, and a normal component obtained as *v*_*n*_ = ***v*** · ***n***, where

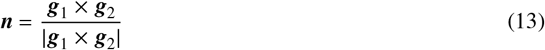

is the unit normal to the surface Γ at ***x***. The second fundamental form of Γ is a second-order symmetric tensor characterizing curvature and given by

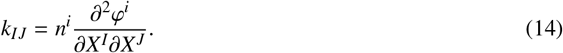

The rate of deformation tensor is given by [44, 56]

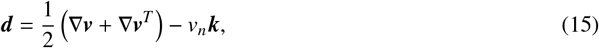

where ∇***v*** is the covariant derivative of ***v*** on Γ.

At each time instant, the derivative of the deformation mapping, *Dφ*(***X***, *t*), is a linear transformation that maps the tangent plane to Γ_0_ (identified with R^2^) at point ***X*** onto the tangent plane to Γ at point *φ*(***X***, *t*), and is often referred to as the deformation gradient denoted by ***F***(***X***, *t*). It describes the deformation of an infinitesimal neighborhood around ***X***. The deformation gradient can be expressed in the natural bases as ***F*** = ***g***_1_ ⊗ ***G***^1^ + ***g***_2_ ⊗ ***G***^2^, with the information about deformation being contained in the natural basis vectors on Γ.

The deformation gradient contains information about local strain and about local rotations. To retain only information about local deformation, we introduce the right Cauchy-Green tensor as [44]

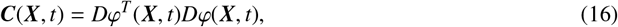

which using the expression of the deformation gradient in terms of the natural bases, can be expressed as

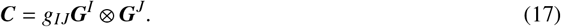

Hence, the components of ***C*** coincide with those of ***g*** = *g*_*IJ*_***g***^*I*^ ⊗ ***g***^*J*^ in the corresponding bases. The Jacobian of the mapping *φ* is defined as

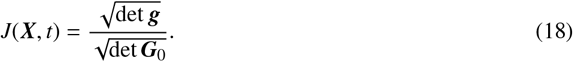

It measures local area changes. Because in Cartesian coordinates det ***G***_0_ = 1, we can write 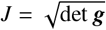.

### A.3 Cortical density and turnover

The areal mass density of the active gel on Γ is denoted by *ρ*(***x***, *t*). If the gel is assumed to be 3D incompressible, then *ρ* can be interpreted as the thickness of the cortical layer. Because *J* represents a ratio between reference and deformed local areas, we can express the mass density per unit reference area as

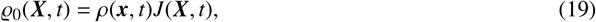

where ***x*** = *φ*(***X***, *t*).

The areal density or thickness of the cortical layer is set by a continuous turnover process involving treadmilling of actin filaments through polymerization and depolymerization of actin monomer or filaments subunits. Here, we use a conventional minimal model for cortical turnover expressed as a continuity equation for *ρ*(***x***, *t*) involving source terms due to polymerization and depolymerization reactions [32, 30, 56]

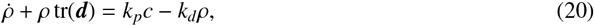

where the dot denotes the material time derivative (at fixed ***X***), and the trace of the rate-of-deformation tensor, measuring the rate of change of local area, can be expressed on a deforming surface as tr(***d***) = ∇ · ***v*** − *v*_*n*_*H*, see Eq. (15) [56]. In the source terms of Eq. (20), *k*_*p*_ is the polymerization rate, *c* is the bulk cytosolic concentration of the cortical material and *k*_*d*_ is the depolymerization rate. Depolymerization is assumed to be proportional to *ρ* as actin filaments can be depolymerized across the thickness whereas polymerization is localized mostly at the cell membrane where the nucleators are present. For a constant pool of cytosolic actin *c* = *c*_0_, i.e. assuming infinite supply of actin monomers, the steady-state cortical thickness is given by

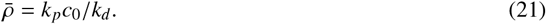

The characteristic timescale for turnover is *τ*_to_ = 1/*k*_*d*_. The turnover equation can be expressed in terms of the reference density *ϱ*_0_ by combining Eqs. (19,20) to obtain the second expression in Eq. (1).

### A.4 Elastic energy with respect to an evolving material metric

Here, we model the short-term nonlinear elasticity of the network of semi-flexible actin filaments using the thermodynamically consistent framework of hyperelasticity [45, 44]. However, to account for the dynamical remodeling of the actin gel, where cross-linkers are transient and filaments turnover, we will introduce the notion of a time-evolving reference configuration as the material is being dynamically rebuilt. To capture this effect, we view the reference metric as a dynamical variable, similar to previous models of nonlinear anelasticity including viscoelasticity, growth or plasticity [46, 47, 48, 49]. In practice, we choose to consider as dynamical variables the contravariant components of the material metric tensor, *G*^*IJ*^(***X***, *t*), whose initial conditions are 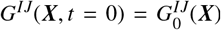. Within this modeling framework, the reference configuration can be understood as a Riemannian manifold with time-evolving metric (Γ_0_, ***G***). Unlike ***G***_0_, ***G*** is not Euclidean, and its incompatibility can result in residual stresses.

The introduction of a time-dependent material metric has a profound impact on the quantification of strain, which in essence is a comparison between the deformation-induced metric tensor ***C*** and the material metric tensor, and for this reason the free-energy density per unit mass of a nonlinearly elastic material is conventionally viewed as a function of both of these tensors, Ψ(***C, G***_0_). For an isotropic material, constitutive models are expressed in terms of the principal invariants of ***C*** relative to ***G***_0_. For a quasi-2D material such as the cortical layer, only two principal invariants are sufficient, for instance the trace 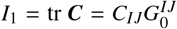 and the determinant 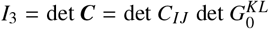 [44], which when using Cartesian coordinates in Γ_0_ simplify to *I*_3_ = *C*_*II*_ and *I*_3_ = det *C*_*IJ*_. Often, the elastic potential Ψ(*I*_1_, *I*_3_) is minimal when the deformation map is the identity map, i.e. when *I*_1_ = 2 and *I*_3_ = 1.

In the present setting, we model the remodeling of the material by comparing ***C***(***X***, *t*) with the time-dependent material metric ***G***(***X***, *t*) in the elastic potential, Ψ(***C, G***). Accordingly, we introduce the invariants relative to the time-dependent material metric as

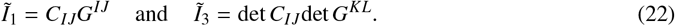

For an isotropic material, the elastic energy per unit mass is 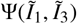. For a finite local deformation given by the symmetric and positive-definite tensor ***C***, now the system can relax the elastic energy by adapting the material metric such that *G*^*IJ*^ = *C*^−1,*IJ*^, since in this case we have 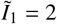 and 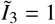. Hence, this formulation can be understood as a tensorial generalization of the notion of dynamical resting length for a 1D spring [3].

The total elastic energy stored in the network can be expressed as

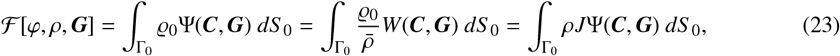

where, to write the last expression, we have used Eq. (19). We have introduced for later use an elastic potential per unit reference area at the steady-state mass density, 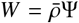. We compute the rate of change of the free energy as

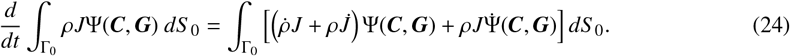

Noting that 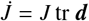 and plugging Eq. (20), we obtain

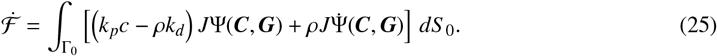

The first term in Eq. (25) accounts for changes in free energy due to changes of mass as a result of turnover, whereas the second term captures changes in elastic energy density due to deformation and evolution of the reference metric. The first term can be seen as a thermodynamic force driving turnover. Here, however, we neglect this effect and assume that active terms driving turnover dominate, leading to Eq. (20). Since this term does not depend on our process mechanical variables, we can ignore it when forming the Rayleighian of the system. Retaining only the second term in Eq. (25) and expanding it, we obtain

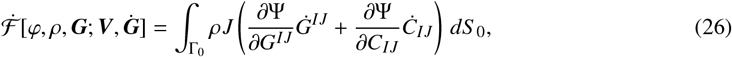

where in the first term, the power conjugate to the rate of change of metric, ∂_***G***_Ψ, is a stress-like quantity driving the evolution of the reference metric. The second term is the usual power of elastic stresses expressed in a fully Lagrangian way,

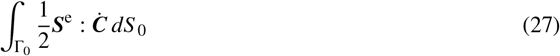

with

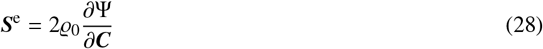

the elastic second Piola-Kirchhoff stress tensor. The elastic power can be expressed in the spatial configuration following a standard calculation as

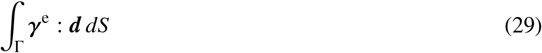

where *γ*^e^ is the elastic part of the Cauchy stress of the thin membrane given by [44]

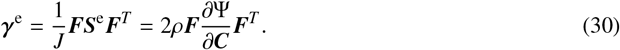

Several strain energy density functions have been proposed to describe the nonlinear elasticity networks of semi-flexible filaments such as actin networks [50]. For the sake of simplicity, we chose an isotropic two-dimensional Neo-Hookean strain energy density given as [45]. While the network may be 3D incompressible, it is 2D compressible [30], and accordingly, we choose the following compressible model

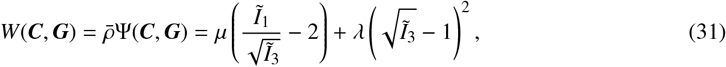

where *λ* and *µ* are 2D Lamé parameters (with units of surface tension) at the steady-state density 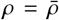, which we assume to be equal [30].

### A.5 Dissipation

The dissipation potential associated with cortical remodeling processes is postulated to be a quadratic potential of the form

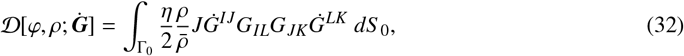

where *η* is a remodeling viscosity coefficient with units of surface viscosity and 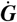 is the material time derivative of the reference metric. See also [87]. As for the standard metric tensors, the covariant and contravariant components of the material metric tensor are related by the relations 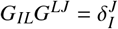.

### A.6 Active power input

We assume that active cortical tension depends on cortical density *γ*^a^(*ρ*) and that the actin filaments in the network are isotropically oriented, which results in an isotropic active cortical tension power-conjugate to the rate of change of local area. Therefore the active power input can be expressed as

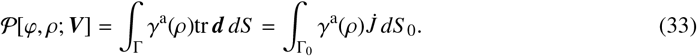

For simplicity, we further follow the common assumption that the active tension is proportional to cortical thickness, *γ*^a^(*ρ*) = *ρξ*, where *ξ* is an activity parameter, although it is known that this relation is more complex and may depend on a more detailed description of network architecture [41, 37].

### A.7 Governing equations of the active viscoelastic cortical behavior

In Onsager’s formalism, the governing equations for the active gel, encoding balance of linear momentum (in tangential and normal directions) and the remodeling evolution law for ***G*** result from the minimization of the Rayleighian defined as

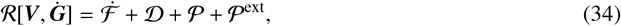

with respect to the process variables ***V*** and 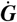. In Eq. (34), P^ext^ accounts for the power supplied by the external tangential tractions ***t*** acting on a subset of the surface boundary ∂Γ_*N*_ ⊂ ∂Γ and possibly a pressure difference *P* across the surface

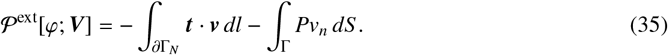

Taking the variation of the Rayleighian with respect to 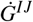 leads to the stationarity condition

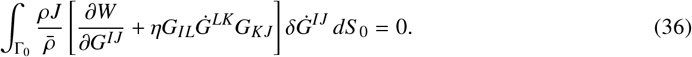

Since the variations 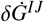 are abitrary, we obtain the following kinetic law for the reference metric tensor

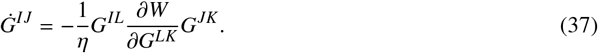

Because for an isotropic material *W* depends on the invariants in Eq. (22), an explicit calculation shows that

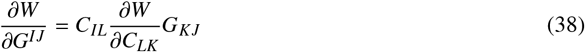

and hence the evolution Eq. (37) can be expressed in terms of the elastic part of the second Piola-Kirchhoff stress tensor ***S***^e^ introduced earlier as

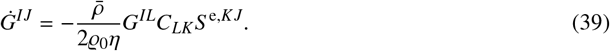

These calculations show that the elastic stresses are the driving force for network remodeling. Because for our Neohookean model ***S***^e^ vanishes whenever *G*^*IJ*^ = *C*^−1,*IJ*^, this equation also shows that elastic stresses dissipate and remodeling ceases when the material metric has relaxed to the current deformation.

The variation of the Rayleighian functional with respect to the tangential surface velocity field ***v*** gives the following stationarity condition

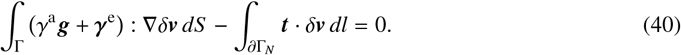

Lastly, the variation of the Rayleighian functional with respect to the normal velocity *v*_*n*_ leads to

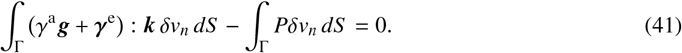

Using the arbitrariness of the variations *δ****v*** and *δv*_*n*_ and integration by parts, we obtain the strong form of momentum balance in the tangential and normal direction to the surface Γ given as

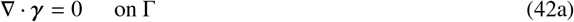

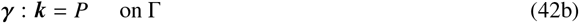

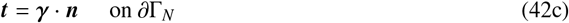

where the total cortical stress (with units of surface tension) includes an active and an elastic contribution, *γ* = *γ*^a^ ***g*** + *γ*^e^. The momentum balance equation in the normal direction is just a generalized form of the Young-Laplace relation, relating the local curvature, surface tension, and pressure difference across the surface. Thus, the dynamics of the cortical active gel surface is governed by balance of linear momentum expressed by Eqs. (42a, 42b, 42c), the evolution law for ***G*** given by Eq. (37), and the statement of mass balance of cortical material in Eq. (20). These equations, independent of the choice of reference configuration, are a system of partial differential equations for the shape, flow, and cortical density of the active gel surface, and a system of ordinary differential equations at each material point for the viscoelastic relaxation.

## B Explicit model for an idealized tissue of hexagonal cells under stretch

We consider here an idealized epithelial tissue made of repeating units of polyhedral cells with flat regular hexagonal apico-basal faces and rectangular lateral faces in the reference state. We further consider two simple loading scenarios on such crystalline tissue: equibiaxial stretching where cells retain their regular hexagonal shape throughout the deformation, and uniaxial stretching along the hexagon’s long axis keeping the width fixed, Fig. 7. In both cases, we assume affinity, i.e. that the deformation of each cell follows that of the tissue. Consequently, it is sufficient to study the dynamics of a single cell.

**Figure 7:**
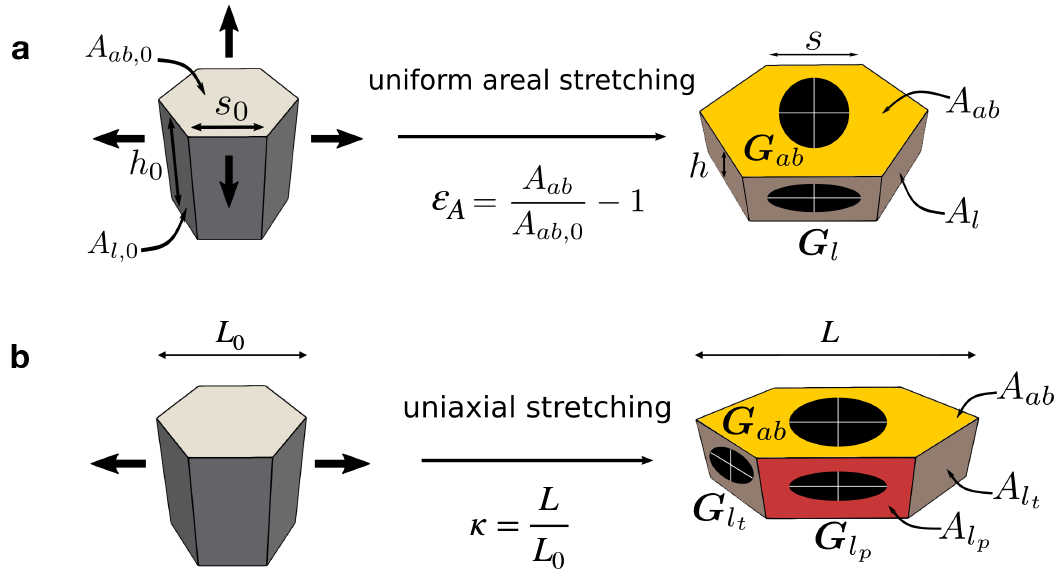
(a) Illustration of the equibiaxial deformation of a cell in an idealized crystalline tissue. The ellipses represent the material metric tensor of apico-basal and of lateral surfaces after relaxation, when they adjust to the imposed deformation. (b) Analogous illustration for a uniaxial deformation.

### B.1 Dynamics under equibiaxial deformation

For a regular hexagonal prism under equibiaxial deformation, all the lateral surfaces are mechanically equivalent. Apical and basal surfaces are also equivalent. We can express the current apico-basal area of the cell as *A*_*ab*_ = (1 + *ε*_*A*_)*A*_*ab*,0_, where *ε*_*A*_ is the tissue areal strain and *A*_*ab*,0_ is the initial apico-basal area of the cell, Fig. 7a. Furthermore, volume conservation, *A*_*ab*,0_*h*_0_ = *A*_*ab*_*h*, allows us to compute the cell height as a function of *ε*_*A*_ as *h* = *V*_0_/*A*_*ab*_ = *h*_0_/(1 + *ε*_*A*_). The surface area of each of the six rectangular lateral faces after deformation, *A*_*l*_ = *sh*, can be computed by noting that the edge of the hexagon deforms according to 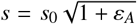, from which we conclude that 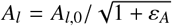. The Jacobians of apico-basal and lateral surfaces are given by

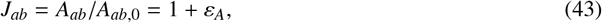

and

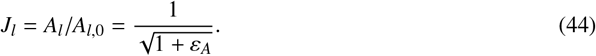

The right Cauchy-Green deformation tensor on the apico-basal faces is isotropic and given by

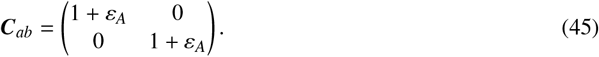

For lateral faces, aligning the Cartesian coordinates with the sides of the rectangle, Fig. 7a, we obtain

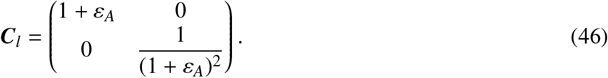

Thus, in such an idealized model, the areal strain of the tissue *ε*_*A*_ parametrizes all kinematic variables. Analogously, by the symmetry of our system, ***G***_*ab*_ is an isotropic tensor, which depends on a single scalar that we denote as *G*_*ab*_, and ***G***_*l*_ is a diagonal tensor, whose diagonal elements are denoted by 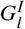 and 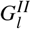. We can thus particularize the elastic potential *W*(***C, G***) to apico-basal and lateral surfaces and express it as functions of our reduced unknowns, that is *W*_*ab*_(*ε*_*A*_, *G*_*ab*_) and 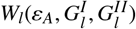. This is clear by recalling Eq. (31) and noting that, on apico-basal faces

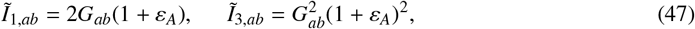

whereas on lateral faces

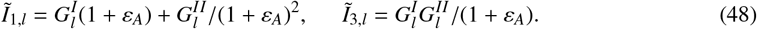

From Eqs. (43,44), we immediately obtain that

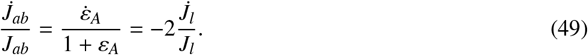

With these expressions and recalling Eq. (1), we obtain the evolution equations for the apico-basal and lateral densities as the ordinary differential equations

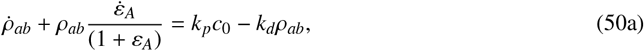

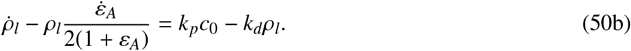

In the unit cell of the crystalline tissue, there are 2 apicobasal surfaces and 6 lateral surfaces. Therefore, the rate of change of free energy in the unit cell is

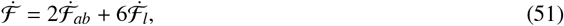

where adapting Eq. (26) to the present setting, we have

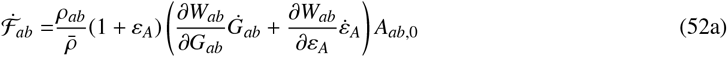

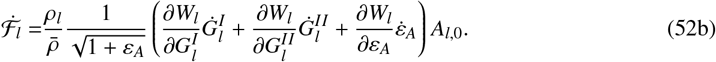

Analogously, we can write the dissipation potential in the unit cell as

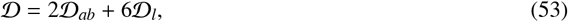

where

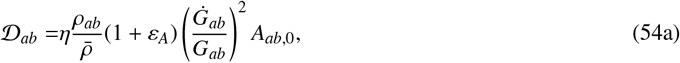

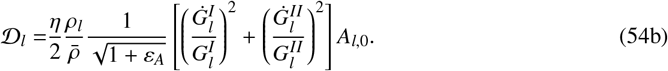

Adapting to the present setting Eq. (33) for the active power input, we obtain

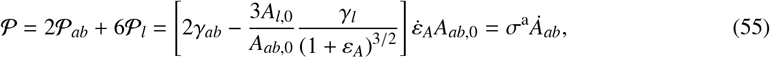

where *γ*_*ab*_ = *ρ*_*ab*_*ξ*_*ab*_ and *γ*_*l*_ = *ρ*_*l*_*ξ*_*l*_ are the apico-basal and lateral active surface tensions. This expression allows us to identify the term between brackets as the active component of tissue tension

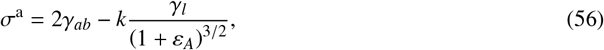

with *k* = 3*A*_*l*,0_/*A*_*ab*,0_ is a fixed geometric constant given by the reference state of the tissue.

To include the power exerted by an externally applied tension *σ* on the tissue, we further include the external power input

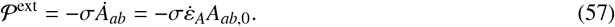

Gathering all the contributions, we write the Rayleighian for the unit cell of the tissue as

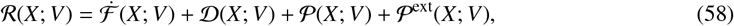

where 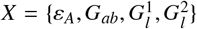 collects all the state variables and *V* the corresponding time derivatives. Minimization of the Rayleighian with respect to the time derivatives of the material tensor components yields

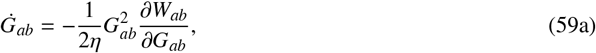

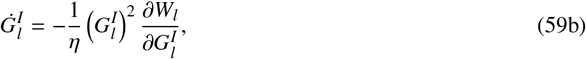

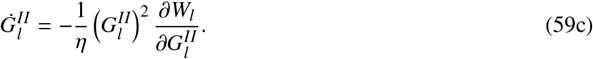

Then, minimizing the Rayleighian with respect to 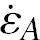, we obtain the expression for the total tissue tension as

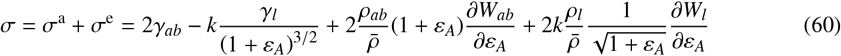

which clearly shows the additive decomposition of tissue tension into its active and its elastic components.

In a scenario where the total tension *σ*(*t*) is given as a function of time, then Eqs. (50,59,60) provide a system of coupled and nonlinear ordinary differential-algebraic equations to solve for 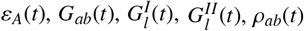and *ρ*_*l*_(*t*). If instead *ε*_*A*_(*t*) is given, then Eq. (60) can be interpreted as a constitutive equation for the tissue, whose evaluation requires the time integration of Eqs. (50,59) to solve for the material metric tensor components and the densities.

For the stress-relaxation scenario studied in Fig. 2b, we compared this idealized model, where we simply need to integrate a small system of differential-algebraic equations, with various tissue models of varying complexity, finding a remarkably good agreement, Fig. 8a. In the reduced tissue model, following a step change in *ε*_*A*_, the relaxation of the densities is exponential, and therefore the relaxation of the active tensions is also exponential. Instead, the relaxation of the material metric tensors, and hence that of tension, is not exponential and can be expressed in terms of Lambert *W* functions. However, the resulting dynamics for *σ*^e^ closely follow an exponential, Fig. 8b. The combined effect of active and elastic relaxations also results in a nearly exponential relaxation of the total tissue tension.

**Figure 8:**
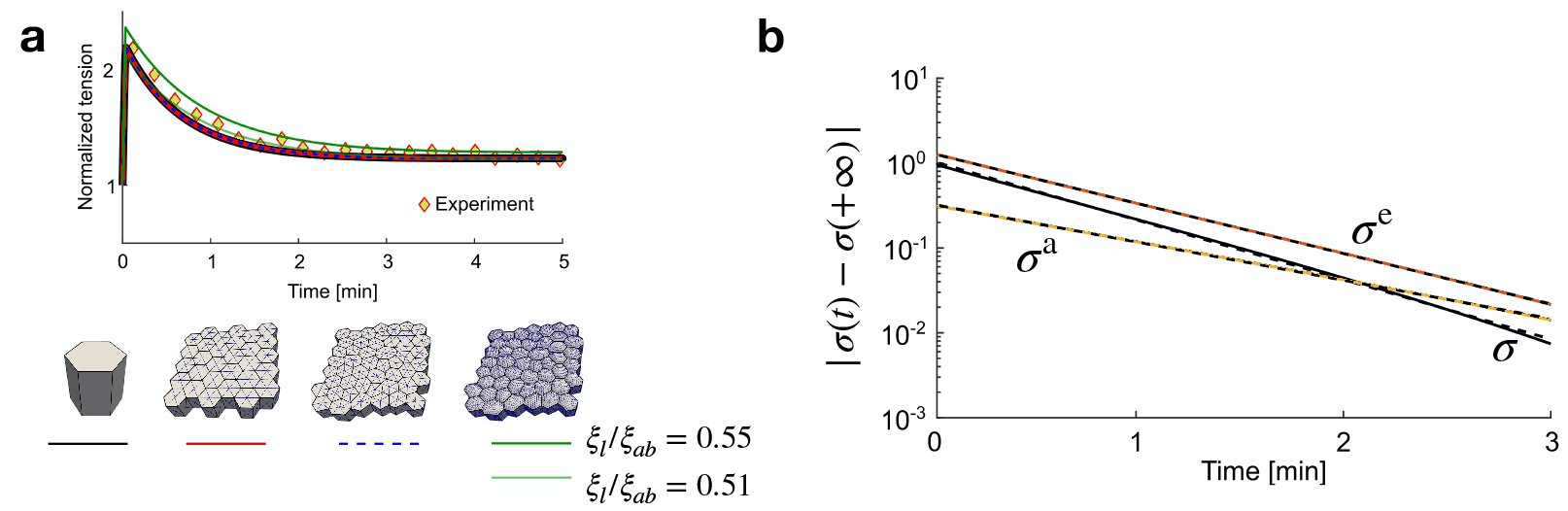
(a) Quantitative agreement in a stress-relaxation scenario (that of Fig. 2b) between the idealized tissue model, a multi-cellular tissue made of hexagonal prismatic cells with the CagVM, and a disordered multicellular tissue with apical cell geometries taken from an experiment with the CagVM and the agVM models. In the agVM model, because of the curvature of cellular faces, a better agreement is found by adjusting the ratio between lateral and apicobasal active tensions. Experimental data are taken from [15]. (b) Total tissue tension (black) along with its elastic (orange) and active (yellow) components as a function of time in semi-logarithmic scale to highlight exponential relaxation. Dashed lines show the exponential fit.

### B.2 Linear stability analysis

To analyze the stability of equibiaxial steady-states of the system, we reformulate our problem by splitting the unknowns between those governed by ordinary differential equations (ODEs), 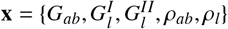, and that governed by an algebraic equation, *y* = *ε*_*A*_. Then, the governing equations can be cast into a system of semi-explicit Differential-Algebraic equations (DAEs) of index-1 [92] of the form

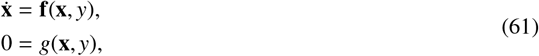

where the first equation encodes Eqs. (50,59) and the second equation encodes Eq. (60). Since *g*(**x**, *y*) = 0 and ∂_*y*_*g* is non-singular, we can use the implicit function theorem to express *y* as a function of **x** and obtain a reduced system of ODEs given by [92]

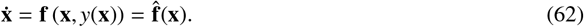

The fixed points **x**^∗^ of this system, characterized by 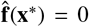, coincide with those of the original DAEs. Linearizing the reduced system about a fixed point **x**^∗^ as **x**(*t*) = **x**^∗^ + *ϵ***z**(*t*), we obtain the linear system of ODEs 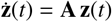, where the matrix **A** is given by

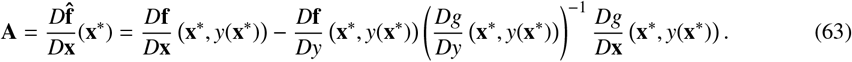

Because the solutions of the linearized system take the form **z**(*t*) = **z**_0_ exp (*t***A**), the stability of the fixed point **x**^∗^ is determined by the spectrum of **A**, which depends on model parameters. The fixed point is stable (unstable) if all (any of the) eigenvalues have negative (positive) real part. The imaginary part of an eigenvalue indicates whether the corresponding stable (unstable) mode evolves towards zero (infinity) with oscillations.

To identify the fixed points of the system, we note that at steady-state *σ* = *σ*^a^, and therefore we can obtain the steady-state strain as

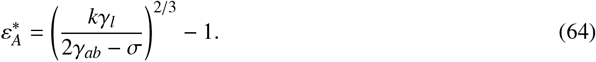

The condition that elastic tension has fully relaxed is met when 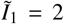 and 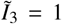 on apicobasal and on lateral surfaces, which using Eqs. (47,48) allows us to identify the steady-state components of the material metric as

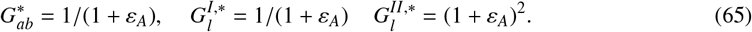

Finally, the steady-state densities are simply 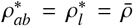. We used Matlab’s symbolic toolbox [93] to evaluate the Jacobian of the system **A** and its spectrum, allowing us to classify the stability of the system across the parameter space. We color-coded steady states with dark green for real positive eigenvalues, light green for complex eigenvalues with positive real part, orange for complex eigenvalues with negative real part and red of real negative eigenvalues, see Fig. 6.

### B.3 Dynamics under uniaxial deformation

Similarly to the equibiaxial case, we derive the kinematics for a cell under uniaxial deformation. We assume that the cell is stretched uniaxially along its long diagonal with a stretch ratio *κ*, Fig. 7b. Under such assumption, the apico-basal surface area changes as *A*_*ab*_ = *κA*_*ab*,0_, with the Jacobian of the deformation on the apico-basal surfaces given by *J*_*ab*_ = *κ* and the right Cauchy-Green deformation tensor given by

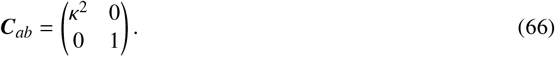

For the lateral surfaces, we need to distinguish the deformation of the two faces parallel to the direction of stretching, which we will refer to 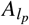, and the deformation of the four other transverse surfaces which we refer to as 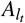, Fig. 7b. Using simple geometric arguments, we can compute the stretch ratios along each edge on a lateral face. Edges aligned in the direction of the applied deformation will be stretched by *κ*. Due to volume conservation of the cell, edges along the height of the cell will be stretched by a ratio of 1/*κ*. Finally, edges in the transverse direction will be stretched by a ratio of 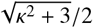. Therefore, the Jacobians of the deformation on the lateral surfaces are given as

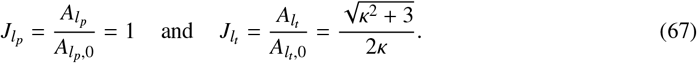

The corresponding Cauchy-Green deformation tensors are given by

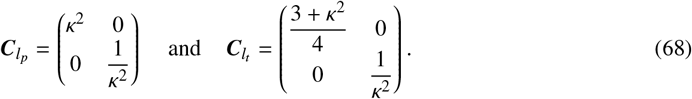

Using the symmetries of the uniaxial setting, now ***G***_*ab*_ is diagonal with two components, along and perpendicular to the stretch direction, which we denote by 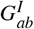and 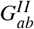. Analogously, the material metric tensor in parallel lateral and transverse lateral surfaces can be described by two scalars along the principal directions. Following an analogous procedure to the equibiaxial one, we can derive ordinary differential equations for the unknowns 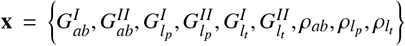, and an algebraic mechanical equation for the uniaxial stretch *y* = *κ*.

### B.4 Modeling actin depletion under stretch

As discussed in [4], epithelial softening can take place as a result of the exhaustion of cytosolic material for polymerization of the actin cortex. To model this in our superelasticity simulations, we assume that the total mass of actin material *M* is constant in each cell, which can be expressed as

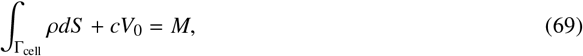

where Γ_cell_ is the surface of a cell and *V*_0_ is its fixed volume. We denote by *S* the total area of the cell and *S* _0_ its total area in a reference configuration. As a cell becomes stretched, *S* increases but *V*_0_ remains constant. In steady-state, combining this equation with the turnover equation, we obtain

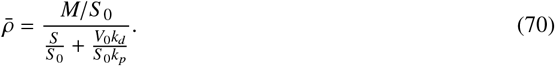

This expression shows that the dimensionless number *V*_0_*k*_*d*_/(*S* _0_*k*_*p*_) controls the sensitivity to stretch-induced softening.

## C Mechanical barrier to large stretch or compression

As argued in the main text, at large stretches, the actomyosin cytoskeleton is thought to cease being the main cytoskeletal structure responsible for tissue tension, as intermediate filaments become engaged. Besides intermediate filaments, the packing of adhesion molecules in shrinking cell-cell junctions or the nucleus trapped between tense apico-basal cortices can also result in strain-stiffening of the tissue. Under compression, crowding at apico-basal surfaces or the nucleus can also result in strain-stiffening.

Even though the strain-stiffening response of intermediate filaments is rate-dependent [19], turnover and re-organizations of these and other structural elements of the cell are much slower than those of the actomyosin cortex, and hence are viewed here as rate-independent components described by an effective free-energy. To account for all these strain-stiffening mechanisms at large compression or stretch in a simplified manner, we consider a rather unspecific mechanical barrier limiting excessive stretching or lateral compression of cells. In our model, this barrier only becomes active at relatively large strains. We implemented a cubic energy functional on each surface that limits individual surface expansion after a certain threshold *J*_high_

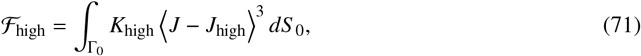

where ⟨ *x*⟩ = *x* if *x >* 0 and 0 otherwise, and *K*_high_ modulates the strength of the resistance to deformations. We also implemented a cubic functional serving as a barrier to surface compression after a certain threshold *J*_low_ of the form

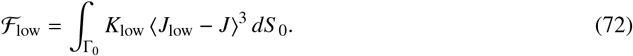

## D Material parameters

Parameter values were estimated from experimental data available in the literature. For instance, the monolayers under consideration have a thickness of about 10 µm–15 µm [13]. Assuming a typical cell edge length *s*_0_ to be around 5 µm [13], we chose the aspect ratio of cells at initial state to be *h*_0_/*s*_0_ = 2. Furthermore, we assumed that apical and basal tensions are contractile, and that contractility on the lateral faces dominates over adhesion so that *ξ*_*a*_, *ξ*_*b*_ and *ξ*_*l*_ are all positive quantities [94]. Due to adhesion energy at cell-cell contacts lowering the lateral surface tension [52], we reasoned that surface tension on lateral cell faces is lower than the average apico-basal surface tension. This ratio *ξ*_*l*_/*ξ*_*ab*_ also determines the steady-state tension after stretch and the contact angle between faces, and was chosen in agreement with the observed geometry of cells resulting in *ξ*_*l*_/*ξ*_*ab*_ ∼ 0.1 − 1. Cortical viscoelastic remodelling due to cross-linker turnover is expected to be faster than the actin turnover such that ∼*τ*_ve_/*τ*_to_ ∼ 0.01 − 0.75 [95, 36, 3, 96]. Following these references, we estimated cortical tensions in the mN/m range and turnover time-scales of tens of seconds. For the ratio between cortical elasticity and active tension, 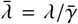, we turned to single cell data in the low frequency limit to estimate that this parameter should be close to one [96]. Finally, the spring-pot material parameters {*C*_*α*_, *α*} were chosen to fit stress relaxation experiments in [3], and are compatible with physical estimates found in the literature [97, 63]. Parameter values for each simulation are summarized in Tables 1–6.

## E Power-law rheology: Materials with memory

For simplicity, we implemented the power-law rheology in a uniaxially stretched idealized tissue, see Appendix B.3. The key step for our modelling framework based on Onsager’s variational formalism is to identify a meaningful rate of change of free energy and dissipation potential for the spring-pot element, which can then be added to the Rayleighian functional. Several works have investigated the formulation of thermodynamically consistent free energy and dissipation functionals for such fractional hereditary materials [98, 99]. For a 1D spring-pot element, the free energy per unit volume is given by

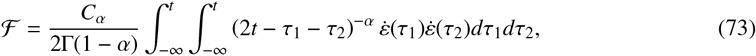

where *C*_*α*_ (with units of Pa · s^*α*^) and *α* are the material parameters characterizing the spring-pot element, and Γ denotes the Gamma function [63]. The free energy in Eq. (73) is fundamentally paired to a specific dissipation rate. By an argument involving the Clausius-Duhem inequality, the associated dissipation rate reads [99]

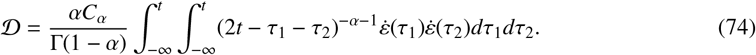

Since the precise cellular structure responsible for the required broad spectrum of relaxation mechanisms is not clear, we associated the spring-pot power-law element to the bulk cytosolic/cytoskeletal shear rate. For a volume-preserving uniaxial deformation of the cytosol, the deformation gradient can be expressed as

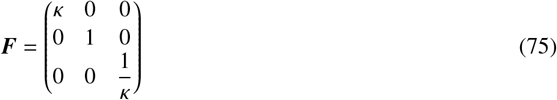

where *κ* is the uniaxial stretch ratio in the tissue, see Appendix B.3. The rate-of-deformation tensor can then be computed as

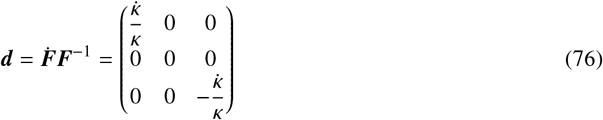

from which we can identify a scalar strain-rate as 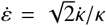. Replacing this Expression in Eq. (73) and multiplying by the cell volume, we obtain

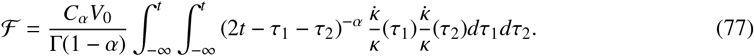

Using the Liebnitz integral rule, we can compute

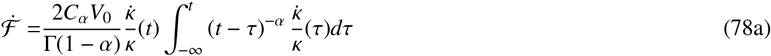

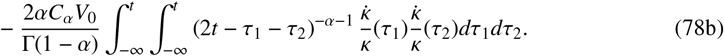

The corresponding dissipation potential is given by

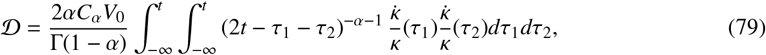

so that it cancels with the last term in Eq. (78b). The effective uniaxial tension due to this effect, *σ*_cyt_, is obtained by taking the variation with respect to the uniaxial strain-rate of the tissue 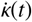

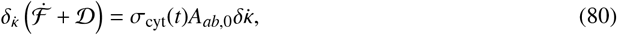

where

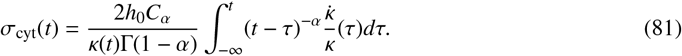

The hereditary integral in this expression for the power-law contribution to the tissue tension is numerically computed using the composite trapezoidal rule.

## Movie captions

**Movie 1:** 10% equibiaxial stretch-unstretch of the CagVM. Because of stretch, cortical density decreases at apico-basal surfaces and increases at lateral surfaces, but due to turnover, cortical density recovers the steady-state over time. The ellipses represent the material metric tensor or generalized resting length in each surface. The dynamic evolution of these ellipses illustrates the viscoelastic relaxation as the material metric tensor tends to the applied deformation. Tension dynamics in the tissue is illustrated with the red arrows. Time is normalized by the turnover timescale.

**Movie 2:** Superelastic response to stretching-unstretching, corresponding to Fig. 2e.

**Movie 3:** Transient tissue buckling after a fast compression, corresponding to Fig. 4a.

**Movie 4:** Different pulsation modes of an unconfined tissue under isotropic and constant tension with high (left) and low (right) cortical elasticity, corresponding to Fig. 6f. Time is normalized by the turnover timescale.

**Movie 5:** Different pulsation modes of a periodic tissue corresponding to Fig. 6g. Time is normalized by the turnover timescale.

**Movie 6:** Self-organized locomotion of a tissue interacting with a substrate by adhesion and friction, corresponding to Fig. 6i,j. Time is normalized by the turnover timescale.

## Notes

### Competing Interest Statement

The authors have declared no competing interest.

### Summary of Updates

Minor edits in the text with some additional references to better reflect the state-of-the-art. Minor changes in the figures and movies (e.g. colormaps for clarity)

## References

[1] Waters, C. M., Roan, E., and Navajas, D. Mechanobiology in lung epithelial cells: Measurements, perturbations, and responses. Comprehensive Physiology, 2:1–29, 2012.

[2] Wong, C. C., Loewke, K. E., Bossert, N. L., Behr, B., De Jonge, C. J., Baer, T. M., and Pera, R. A. R. Non-invasive imaging of human embryos before embryonic genome activation predicts development to the blastocyst stage. Nature Biotechnology, 28:1115–1121, 2010.

[3] Khalilgharibi, N., et al. Stress relaxation in epithelial monolayers is controlled by the actomyosin cortex. Nature Physics, 15:839–847, 2019.

[4] Latorre, E., et al. Active superelasticity in three-dimensional epithelia of controlled shape. Nature, 563:203–208, 2018.

[5] Gudipaty, S. A., Lindblom, J., Loftus, P. D., Redd, M. J., Edes, K., Davey, C. F., Krishnegowda, V., and Rosenblatt, J. Mechanical stretch triggers rapid epithelial cell division through Piezo1. Nature, 543:118–121, 2017.

[6] Guillot, C. and Lecuit, T. Mechanics of Epithelial Tissue Homeostasis and Morphogenesis. Science, 340:1185–1189, 2013.

[7] Solon, J., Kaya-Çopur, A., Colombelli, J., and Brunner, D. Pulsed Forces Timed by a Ratchet-like Mechanism Drive Directed Tissue Movement during Dorsal Closure. Cell, 137:1331–1342, 2009.

[8] Hannezo, E., Prost, J., and Joanny, J.-F. Theory of epithelial sheet morphology in three dimensions. Proceedings of the National Academy of Sciences, 111:27–32, 2014.

[9] Nelson, C. M. On Buckling Morphogenesis. Journal of Biomechanical Engineering, 138, 2016. 021005.

[10] Sawyer, J. M., Harrell, J. R., Shemer, G., Sullivan-Brown, J., Roh-Johnson, M., and Goldstein, B. Apical constriction: A cell shape change that can drive morphogenesis. Developmental Biology, 341:5–19, 2010.

[11] Pérez-González, C., et al. Mechanical compartmentalization of the intestinal organoid enables crypt folding and collective cell migration. Nature Cell Biology, 23:745–757, 2021.

[12] Gómez-González, M., Latorre, E., Arroyo, M., and Trepat, X. Measuring mechanical stress in living tissues. Nature Reviews Physics, 2:300–317, 2020.

[13] Harris, A. R., Peter, L., Bellis, J., Baum, B., Kabla, A. J., and Charras, G. T. Characterizing the mechanics of cultured cell monolayers. Proceedings of the National Academy of Sciences, 109:16449– 16454, 2012.

[14] Harris, A. R., Bellis, J., Khalilgharibi, N., Wyatt, T., Baum, B., Kabla, A. J., and Charras, G. T. Generating suspended cell monolayers for mechanobiological studies. Nature Protocols, 8:2516–2530, 2013.

[15] Casares, L., Vincent, R., Zalvidea, D., Campillo, N., Navajas, D., Arroyo, M., and Trepat, X. Hydraulic fracture during epithelial stretching. Nature Materials, 14:343–351, 2015.

[16] Marín-Llauradó, A., et al. Mapping mechanical stress in curved epithelia of designed size and shape. Nature Communications, 14:4014, 2023.

[17] Wyatt, T. P. J., Fouchard, J., Lisica, A., Khalilgharibi, N., Baum, B., Recho, P., Kabla, A. J., and Charras, G. T. Actomyosin controls planarity and folding of epithelia in response to compression. Nature Materials, 19:109–117, 2020.

[18] Fouchard, J., Wyatt, T. P. J., Proag, A., Lisica, A., Khalilgharibi, N., Recho, P., Suzanne, M., Kabla, A., and Charras, G. Curling of epithelial monolayers reveals coupling between active bending and tissue tension. Proc. Natl. Acad. Sci. U. S. A., 117:9377–9383, 2020.

[19] Duque, J., Bonfanti, A., Fouchard, J., Baldauf, L., Azenha, S. R., Ferber, E., Harris, A., Barriga, E. H., Kabla, A. J., and Charras, G. Rupture strength of living cell monolayers. Nature Materials, 23:1563–1574, 2024.

[20] Nagai, T. and Honda, H. A dynamic cell model for the formation of epithelial tissues. Philosophical Magazine B: Physics of Condensed Matter; Statistical Mechanics, Electronic, Optical and Magnetic Properties, 81:699–719, 2001.

[21] Fletcher, A. G., Osterfield, M., Baker, R. E., and Shvartsman, S. Y. Vertex models of epithelial morphogenesis. Biophysical Journal, 106:2291–2304, 2014.

[22] Alt, S., Ganguly, P., and Salbreux, G. Vertex models: from cell mechanics to tissue morphogenesis. Philosophical Transactions of the Royal Society of London B: Biological Sciences, 372:20150520, 2017.

[23] Bi, D., Yang, X., Marchetti, M. C., and Manning, M. L. Motility-driven glass and jamming transitions in biological tissues. Phys. Rev. X, 6:021011, 2016.

[24] Kim, S., Pochitaloff, M., Stooke-Vaughan, G. A., and Campás, O. Embryonic tissues as active foams. Nature Physics, 17:859–866, 2021.

[25] Runser, S., Vetter, R., and Iber, D. Simucell3d: three-dimensional simulation of tissue mechanics with cell polarization. Nature Computational Science, 4:299–309, 2024.

[26] Muñoz, J. J. and Albo, S. Physiology-based model of cell viscoelasticity. Physical Review E, 88:012708, 2013.

[27] Lenne, P.-F., Rupprecht, J.-F., and Viasnoff, V. Cell junction mechanics beyond the bounds of adhesion and tension. Developmental Cell, 56:202–212, 2021.

[28] Moisdon, E., Seez, P., Molino, F. m. c., Marcq, P., and Gay, C. Mapping cell cortex rheology to tissue rheology and vice versa. Phys. Rev. E, 106:034403, 2022.

[29] Prost, J., Jülicher, F., and Joanny, J.-F. Active gel physics. Nature Physics, 2015.

[30] Salbreux, G., Prost, J., and Joanny, J. F. Hydrodynamics of Cellular Cortical Flows and the Formation of Contractile Rings. Physical Review Letters, 103:058102, 2009.

[31] Callan-Jones, A. C. and Voituriez, R. Active gel model of amoeboid cell motility. New Journal of Physics, 15, 2013.

[32] Turlier, H., Audoly, B., Prost, J., and Joanny, J. F. Furrow constriction in animal cell cytokinesis. Biophysical Journal, 106:114–123, 2014.

[33] Mogilner, A. and Manhart, A. Intracellular fluid mechanics: Coupling cytoplasmic flow with active cytoskeletal gel. Annual Review of Fluid Mechanics, 50:347–370, 2018.

[34] Staddon, M. F., Munro, E. M., and Banerjee, S. Pulsatile contractions and pattern formation in excitable actomyosin cortex. PLOS Computational Biology, 18:1–21, 2022.

[35] Mirza, W., De Corato, M., Pensalfini, M., Vilanova, G., Torres-Sánchez, A., and Arroyo, M. Theory of active self-organization of dense nematic structures in the actin cytoskeleton. eLife, 13:RP93097, 2024.

[36] Salbreux, G., Charras, G., and Paluch, E. Actin cortex mechanics and cellular morphogenesis. Trends in Cell Biology, 22:536–545, 2012.

[37] Koenderink, G. H. and Paluch, E. K. Architecture shapes contractility in actomyosin networks. Current Opinion in Cell Biology, 50:79–85, 2018.

[38] Chugh, P. and Paluch, E. K. The actin cortex at a glance. Journal of cell science, 131:jcs186254, 2018.

[39] Leadbetter, T., Seiphoori, A., Reina, C., and Purohit, P. K. Emergence of viscosity and dissipation via stochastic bonds. Journal of the Mechanics and Physics of Solids, 158:104660, 2022.

[40] Belmonte, J. M., Leptin, M., and Nédélec, F. A theory that predicts behaviors of disordered cytoskeletal networks. Mol Syst Biol, 13, 2017.

[41] Chugh, P., Clark, A. G., Smith, M. B., Cassani, D. A. D., Dierkes, K., Ragab, A., Roux, P. P., Charras, G., Salbreux, G., and Paluch, E. K. Actin cortex architecture regulates cell surface tension. Nature Cell Biology, 2017.

[42] Kruse, K., Joanny, J. F., Jülicher, F., Prost, J., and Sekimoto, K. Generic theory of active polar gels: A paradigm for cytoskeletal dynamics. European Physical Journal E, 16:5–16, 2005.

[43] Banerjee, D. S., Munjal, A., Lecuit, T., and Rao, M. Actomyosin pulsation and flows in an active elastomer with turnover and network remodeling. Nature Communications, 8:1121, 2017.

[44] Marsden, J. E. and Hughes, T. J. R. Mathematical foundations of elasticity. Dover, New York, 1994.

[45] Holzapfel, G. A. Nonlinear solid mechanics: a continuum approach for engineering. Wiley, Chichester ; New York, 2000. ISBN 9780471823049 9780471823193.

[46] Kumar, A. and Lopez-Pamies, O. On the two-potential constitutive modeling of rubber viscoelastic materials. Comptes Rendus Mécanique, 344:102–112, 2016.

[47] Miehe, C. A constitutive frame of elastoplasticity at large strains based on the notion of a plastic metric. International Journal of Solids and Structures, 35:3859–3897, 1998.

[48] Sadik, S. and Yavari, A. Nonlinear anisotropic viscoelasticity. Journal of the Mechanics and Physics of Solids, 182:105461, 2024.

[49] Kumar, A. and Yavari, A. Nonlinear mechanics of remodeling. Journal of the Mechanics and Physics of Solids, 181:105449, 2023.

[50] Meng, F. and Terentjev, E. M. Nonlinear elasticity of semiflexible filament networks. Soft Matter, 12:6749–6756, 2016.

[51] Bernstein, B. and Rajagopal, K. Thermodynamics of hypoelasticity. Zeitschrift für angewandte Mathematik und Physik, 59:537–553, 2008.

[52] Mâitre, J.-L. and Heisenberg, C.-P. Three functions of cadherins in cell adhesion. Current Biology, 23:R626–R633, 2013.

[53] Kaurin, D., Bal, P. K., and Arroyo, M. Peeling dynamics of fluid membranes bridged by molecular bonds: moving or breaking. J. R. Soc. Interface, 19, 2022.

[54] Jiang, H. and Sun, S. X. Cellular pressure and volume regulation and implications for cell mechanics. Biophysical Journal, 105:609–619, 2013.

[55] Cadart, C., Venkova, L., Recho, P., Lagomarsino, M. C., and Piel, M. The physics of cell-size regulation across timescales. Nature Physics, 15:993–1004, 2019.

[56] Torres-Sánchez, A., Millán, D., and Arroyo, M. Modelling fluid deformable surfaces with an emphasis on biological interfaces. Journal of Fluid Mechanics, 872:218–271, 2019.

[57] Ishimoto, Y. and Morishita, Y. Bubbly vertex dynamics: A dynamical and geometrical model for epithelial tissues with curved cell shapes. Phys. Rev. E, 90:052711, 2014.

[58] Mâitre, J.-L., Turlier, H., Illukkumbura, R., Eismann, B., Niwayama, R., Nédélec, F., and Hiiragi, T. Asymmetric division of contractile domains couples cell positioning and fate specification. Nature, 536:344–348, 2016.

[59] Drozdowski, O. M. and Schwarz, U. S. Cell bulging and extrusion in a three-dimensional bubbly vertex model for curved epithelial sheets, 2024. URL https://arxiv.org/abs/2411.07141.

[60] Pensalfini, M., Golde, T., Trepat, X., and Arroyo, M. Nonaffine Mechanics of Entangled Networks Inspired by Intermediate Filaments. Physical Review Letters, 131:058101, 2023.

[61] Miehe, C. Computational micro-to-macro transitions for discretized micro-structures of heterogeneous materials at finite strains based on the minimization of averaged incremental energy. Computer Methods in Applied Mechanics and Engineering, 192:559–591, 2003.

[62] Fischer-Friedrich, E., Toyoda, Y., Cattin, C. J., Müller, D. J., Hyman, A. A., and Jülicher, F. Rheology of the Active Cell Cortex in Mitosis. Biophysical Journal, 111:589–600, 2016.

[63] Bonfanti, A., Fouchard, J., Khalilgharibi, N., Charras, G., and Kabla, A. A unified rheological model for cells and cellularised materials. Royal Society Open Science, 7:190920, 2020.

[64] Choudhury, M. I., Benson, M. A., and Sun, S. X. Trans-epithelial fluid flow and mechanics of epithelial morphogenesis. Seminars in Cell & Developmental Biology, 131:146–159, 2022.

[65] Torres-Sánchez, A., Kerr Winter, M., and Salbreux, G. Tissue hydraulics: Physics of lumen formation and interaction. Cells & Development, 168:203724, 2021. Quantitative Cell and Developmental Biology.

[66] Chahare, N., Ouzeri, A., Wilson, T., Bal, P. K., Golde, T., Vilanova, G., Pujol-Vives, P., Roca-Cusachs, P., Trepat, X., and Arroyo, M. Multiscale wrinkling dynamics in epithelial shells. bioRxiv, 2025.06.30.662426, 2025.

[67] Lisica, A., Fouchard, J., Kelkar, M., Wyatt, T. P. J., Duque, J., Ndiaye, A.-B., Bonfanti, A., Baum, B., Kabla, A. J., and Charras, G. T. Tension at intercellular junctions is necessary for accurate orientation of cell division in the epithelium plane. Proceedings of the National Academy of Sciences, 119:e2201600119, 2022.

[68] Martin, A. C., Kaschube, M., and Wieschaus, E. F. Pulsed contractions of an actin-myosin network drive apical constriction. Nature, 457:495–499, 2009.

[69] Azevedo, D., Antunes, M., Prag, S., Ma, X., Hacker, U., Brodland, G. W., Hutson, M. S., Solon, J., and Jacinto, A. DRhoGEF2 Regulates Cellular Tension and Cell Pulsations in the Amnioserosa during Drosophila Dorsal Closure. PLoS ONE, 6:e23964, 2011.

[70] Armon, S., Bull, M. S., Aranda-Diaz, A., and Prakash, M. Ultrafast epithelial contractions provide insights into contraction speed limits and tissue integrity. Proceedings of the National Academy of Sciences of the United States of America, page 201802934, 2018.

[71] Serra-Picamal, X., Conte, V., Vincent, R., Anon, E., Tambe, D. T., Bazellieres, E., Butler, J. P., Fredberg, J. J., and Trepat, X. Mechanical waves during tissue expansion. Nature Physics, 8:628–634, 2012.

[72] Gross, P., Kumar, K. V., and Grill, S. W. How Active Mechanics and Regulatory Biochemistry Combine to Form Patterns in Development. Annual Review of Biophysics, 2017.

[73] Boocock, D., Hino, N., Ruzickova, N., Hirashima, T., and Hannezo, E. Theory of mechanochemical patterning and optimal migration in cell monolayers. Nature Physics, 17:267–274, 2021.

[74] Munjal, A., Philippe, J.-M., Munro, E., and Lecuit, T. A self-organized biomechanical network drives shape changes during tissue morphogenesis. Nature, 524:351–355, 2015.

[75] Lin, S.-Z., Xue, S.-L., Li, B., and Feng, X.-Q. An oscillating dynamic model of collective cells in a monolayer. Journal of the Mechanics and Physics of Solids, 112:650–666, 2018.

[76] Nishikawa, M., Naganathan, S. R., Jülicher, F., and Grill, S. W. Controlling contractile instabilities in the actomyosin cortex. eLife, 6:e19595, 2017.

[77] Muñoz, J. J., Dingle, M., and Wenzel, M. Mechanical oscillations in biological tissues as a result of delayed rest-length changes. Phys. Rev. E, 98:052409, 2018.

[78] Lin, S.-Z., Li, B., Lan, G., and Feng, X.-Q. Activation and synchronization of the oscillatory mor-phodynamics in multicellular monolayer. Proceedings of the National Academy of Sciences, 114:8157–8162, 2017.

[79] Dierkes, K., Sumi, A., Solon, J., and Salbreux, G. Spontaneous oscillations of elastic contractile materials with turnover. Physical Review Letters, 113:1–5, 2014.

[80] Armon, S., Bull, M. S., Moriel, A., Aharoni, H., and Prakash, M. Modeling epithelial tissues as active-elastic sheets reproduce contraction pulses and predict rip resistance. Communications Physics, 4:216, 2021.

[81] Krajnc, M. and Ziherl, P. Theory of epithelial elasticity. Physical Review E, 92:052713, 2015.

[82] Ishihara, S., Marcq, P., and Sugimura, K. From cells to tissue: A continuum model of epithelial mechanics. Phys. Rev. E, 96:022418, 2017.

[83] Morris, R. G. and Rao, M. Active morphogenesis of epithelial monolayers. Phys. Rev. E, 100:022413, 2019.

[84] Haas, P. A. and Goldstein, R. E. Nonlinear and nonlocal elasticity in coarse-grained differential-tension models of epithelia. Phys. Rev. E, 99:022411, 2019.

[85] Andrenšek, U., Ziherl, P., and Krajnc, M. Wrinkling Instability in Unsupported Epithelial Sheets. Physical Review Letters, 130:198401, 2023.

[86] Drozdowski, O. M. and Schwarz, U. S. Morphological instability at topological defects in a three-dimensional vertex model for spherical epithelia. Phys. Rev. Res., 6:L022045, 2024.

[87] Bal, P. K., Ouzeri, A., and Arroyo, M. Continuum theory for the mechanics of curved epithelial shells by coarse-graining an ensemble of active gel cellular surfaces. bioRxiv, 2025.02.27.640501, 2025.

[88] Gómez-Gálvez, P., et al. Scutoids are a geometrical solution to three-dimensional packing of epithelia. Nature Communications, 9:2960, 2018.

[89] Torres-Sánchez, A., Kerr Winter, M., and Salbreux, G. Interacting active surfaces: A model for three-dimensional cell aggregates. PLOS Computational Biology, 18:1–25, 2022.

[90] Arroyo, M., Walani, N., Torres-Sánchez, A., and Kaurin, D. Onsager’s variational principle in soft matter : introduction and application to the dynamics of adsorption of proteins onto fluid membranes. The role of mechanics in the study of lipid bilayers, pages 1–53, 2017.

[91] Doi, M. Onsager’s variational principle in soft matter. Journal of Physics Condensed Matter, 2011.

[92] Riaza, R. Differential-algebraic systems: analytical aspects and circuit applications. World Scientific, New Jersey, 2008. ISBN 9789812791801. OCLC: ocn198759476.

[93] MATLAB. 9.3.0.713579 (R2017b). The MathWorks Inc., Natick, Massachusetts, 2017.

[94] Maitre, J.-L., Berthoumieux, H., Krens, S. F. G., Salbreux, G., Julicher, F., Paluch, E., and Heisenberg, C.-P. Adhesion Functions in Cell Sorting by Mechanically Coupling the Cortices of Adhering Cells. Science, 338:253–256, 2012.

[95] Murthy, K. and Wadsworth, P. Myosin-II-Dependent Localization and Dynamics of F-Actin during Cytokinesis. Current Biology, 15:724–731, 2005.

[96] Hosseini, K., Sbosny, L., Poser, I., and Fischer-Friedrich, E. Binding Dynamics of α-Actinin-4 in Dependence of Actin Cortex Tension. Biophysical Journal, 119:1091–1107, 2020.

[97] Bonakdar, N., Gerum, R., Kuhn, M., Spörrer, M., Lippert, A., Schneider, W., Aifantis, K. E., and Fabry, B. Mechanical plasticity of cells. Nature Materials, 15:1090–1094, 2016.

[98] Lion, A. On the thermodynamics of fractional damping elements. Continuum Mechanics and Ther-modynamics, 9:83–96, 1997.

[99] Deseri, L., Di Paola, M., and Zingales, M. Free energy and states of fractional-order hereditariness. International Journal of Solids and Structures, 51:3156–3167, 2014.

